# Extracellular Granzyme B Promotes Melanocytorrhagy, Melanocyte Senescence and Aberrant Epidermal Differentiation in Vitiligo and Is Therapeutically Targetable

**DOI:** 10.64898/2026.09.19.752829

**Authors:** Siwei Diao, Lingye Jin, Beáta Szilvia Bolla, Hiroto Kuwabiraki, Yasutaka Kuroda, Fei Yang, Lingli Yang, Ichiro Katayama, Daisuke Tsuruta, David J. Granville, Sho Hiroyasu

## Abstract

**Background:** Vitiligo is an acquired depigmenting disorder characterised by melanocyte loss. While immune responses against melanocytes are implicated in its pathogenesis, melanocyte detachment, melanocyte senescence and aberrant epidermal differentiation are increasingly characterised as additional pathological features. Granzyme B (GzmB), classically recognised as an intracellular effector of perforin-dependent cytotoxicity, is increased in vitiligo lesions; however, its extracellular contribution to vitiligo pathology remains unclear.

**Objectives:** To determine whether extracellular GzmB contributes to depigmentation and epidermal pathological features relevant to vitiligo, and to define its cellular source and underlying mechanisms.

**Methods:** Human vitiligo skin samples, murine depigmentation models and cultured human epidermal cells were analysed. The functional effects of extracellular GzmB were examined by subcutaneous administration of recombinant GzmB in mice. A rhododendrol (RD)-induced leukoderma model was used to evaluate the disease relevance of GzmB and the therapeutic effect of topical GzmB inhibition with VTI-1002.

**Results:** GzmB-positive cells were markedly increased in active vitiligo lesions and were predominantly associated with tryptase-positive cells, with limited co-distribution with CD8 or perforin. Vitiligo lesions also showed melanocyte detachment, increased p16^INK4A^-positive melanocytes and aberrant keratinocyte differentiation. These pathological features were recapitulated in vivo by extracellular GzmB administration, accompanied by focal depigmentation. In cultured melanocytes, GzmB exerted no cytotoxic effects but reduced attachment strength and induced a senescence-associated phenotype characterised by decreased extracellular matrix- and adhesion-related gene expression, increased p16^INK4A^ level, elevated senescence-associated β-galactosidase activity and activation of transforming growth factor-β/SMAD signalling. In keratinocytes, extracellular GzmB promoted aberrant differentiation associated with activation of p53 signalling. Topical inhibition of GzmB in RD-induced leukoderma attenuated depigmentation progression, melanocyte detachment, melanocyte senescence and abnormal epidermal differentiation.

**Conclusions:** Extracellular GzmB promotes depigmentation associated with inducing melanocyte detachment, melanocyte senescence and aberrant keratinocyte differentiation. These findings identify extracellular GzmB as a previously underrecognised pathogenic mediator and potential therapeutic target in vitiligo-associated depigmentation.

**What is already known about this topic?:**

- Vitiligo is an acquired depigmenting disorder characterised by melanocyte loss.
- Although autoreactive immune responses are implicated in vitiligo, melanocyte detachment, melanocyte senescence and keratinocyte abnormalities are increasingly recognised as additional pathological features that may also contribute to depigmentation.
- Granzyme B is increased in vitiligo lesions, but whether extracellular granzyme B contributes to disease pathology independently of classical perforin-dependent cytotoxicity has not been tested.

**What does this study add?:**

- In active vitiligo lesions, granzyme B was increased and predominantly distributed to perforin-negative, tryptase-positive mast cells.
- Extracellular granzyme B induced focal depigmentation, melanocyte detachment, melanocyte senescence and aberrant keratinocyte differentiation.
- Pharmacological inhibition of extracellular granzyme B attenuated depigmentation progression and associated pathological changes in a rhododendrol-induced vitiligo-like mouse model.

**What is the translational message?:**

- Granzyme B contributes to melanocyte loss via extracellular mechanisms beyond classical intracellular cytotoxicity.
- These findings support topical inhibition of extracellular granzyme B as a potential local therapeutic strategy for active vitiligo.

**Abstract figure:** Abbreviated abstract (Teaser Text)
In vitiligo, mast cell-associated extracellular granzyme B promotes depigmentation accompanied by melanocyte detachment, melanocyte senescence and aberrant keratinocyte differentiation. Topical granzyme B inhibition attenuates depigmentation and associated pathological changes in a rhododendrol-induced vitiligo-like mouse model, identifying extracellular granzyme B as a potential therapeutic target for depigmenting skin disease.

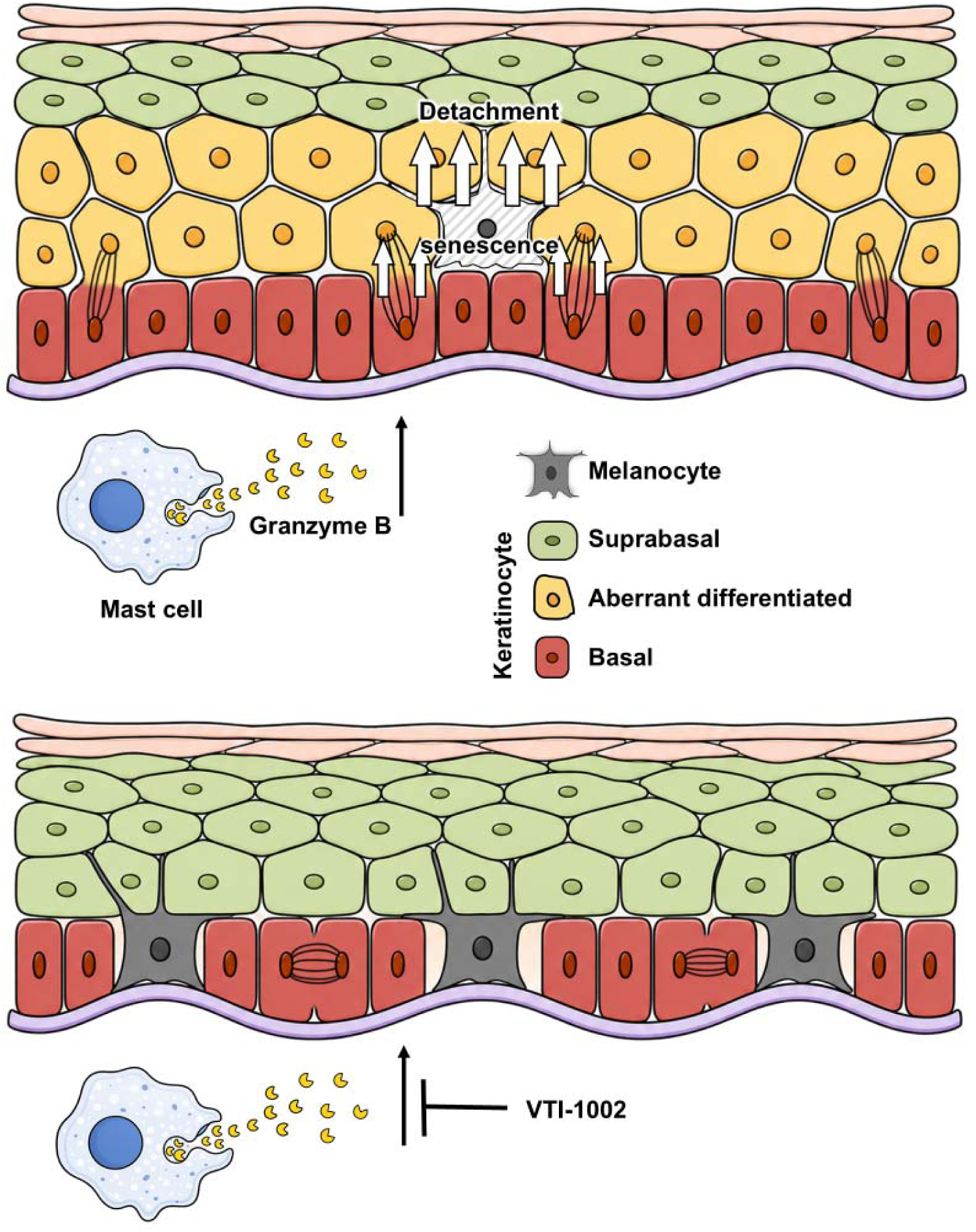

## Introduction

Vitiligo is an acquired depigmenting disorder caused by the loss of epidermal melanocytes and characterised by depigmented macules.^1^ Visible depigmentation imposes a substantial psychosocial burden.^2^ Current treatment options remain limited, with incomplete responses and frequent relapses despite immunomodulators and phototherapy,^3^ highlighting the need for additional therapeutic strategies and a better understanding of the underlying mechanisms.^1,3^

To date, CD8-positive cytotoxic T cell responses against melanocytes have been proposed as key contributors to vitiligo pathogenesis.^1,3–6^ However, accumulating evidence suggests that cytotoxicity alone does not account for the full spectrum of vitiligo pathology.^7^ Beyond immune-mediated melanocyte killing, additional pathological processes have been described, including melanocyte detachment from the basal epidermal layer followed by transepidermal elimination (melanocytorrhagy),^8,9^ melanocyte dysfunction characterised by a senescent phenotype,^10^ and keratinocyte abnormalities involving aberrant differentiation and stress-associated states.^11,12^ Altered keratinocyte division has also been associated with adjacent melanocyte elimination.^13^ Although each of these features has been implicated in melanocyte loss, the upstream mechanisms coordinating these pathological changes remain unclear.

Granzyme B (GzmB) is a serine protease classically recognised as a cytotoxic effector molecule released by CD8-positive T cells and natural killer cells together with perforin.^14^ In this canonical pathway, perforin enables the intracellular delivery of GzmB into target cells, inducing apoptosis during immune responses to infection and tumours.^15,16^ However, beyond this cytotoxic context, recent studies, including ours, have revealed that GzmB is also released by mast cells and other immune cells in a perforin-independent manner and persists in the extracellular space.^17,18^ Extracellular GzmB has been implicated in the pathogenesis of inflammatory skin conditions by affecting extracellular tissue homeostasis, cytokine activity and cellular responses.^19^

In vitiligo, GzmB is increased in both peripheral blood and lesional skin,^20,21^ and *GZMB* polymorphisms are associated with disease susceptibility.^22,23^ However, previous work mainly considered GzmB in the context of intracellular cytotoxicity, and whether extracellular GzmB contributes to vitiligo-associated epidermal phenotypes remains untested.

In this study, we explored the extracellular role of GzmB in vitiligo-associated pathological changes using subcutaneous GzmB administration in mice, a rhododendrol (RD)-induced murine vitiligo-like model, cultured epidermal cells and human skin. Our data demonstrate that extracellular GzmB promotes depigmentation, accompanied by melanocyte detachment, melanocyte senescence and aberrant keratinocyte differentiation, revealing a previously unrecognised extracellular pathogenic function of GzmB in vitiligo.

## Materials and methods

Materials and methods are provided in the Supporting Information.

## Results

### Active vitiligo lesions exhibit melanocyte detachment, melanocyte senescence, aberrant keratinocyte differentiation and increased mast cell-associated granzyme B with limited perforin association

To characterise epidermal pathological changes in active vitiligo, we analysed lesions that had newly developed or enlarged within the previous 6 months (Figure 1a, Table 1).

**Figure 1.**
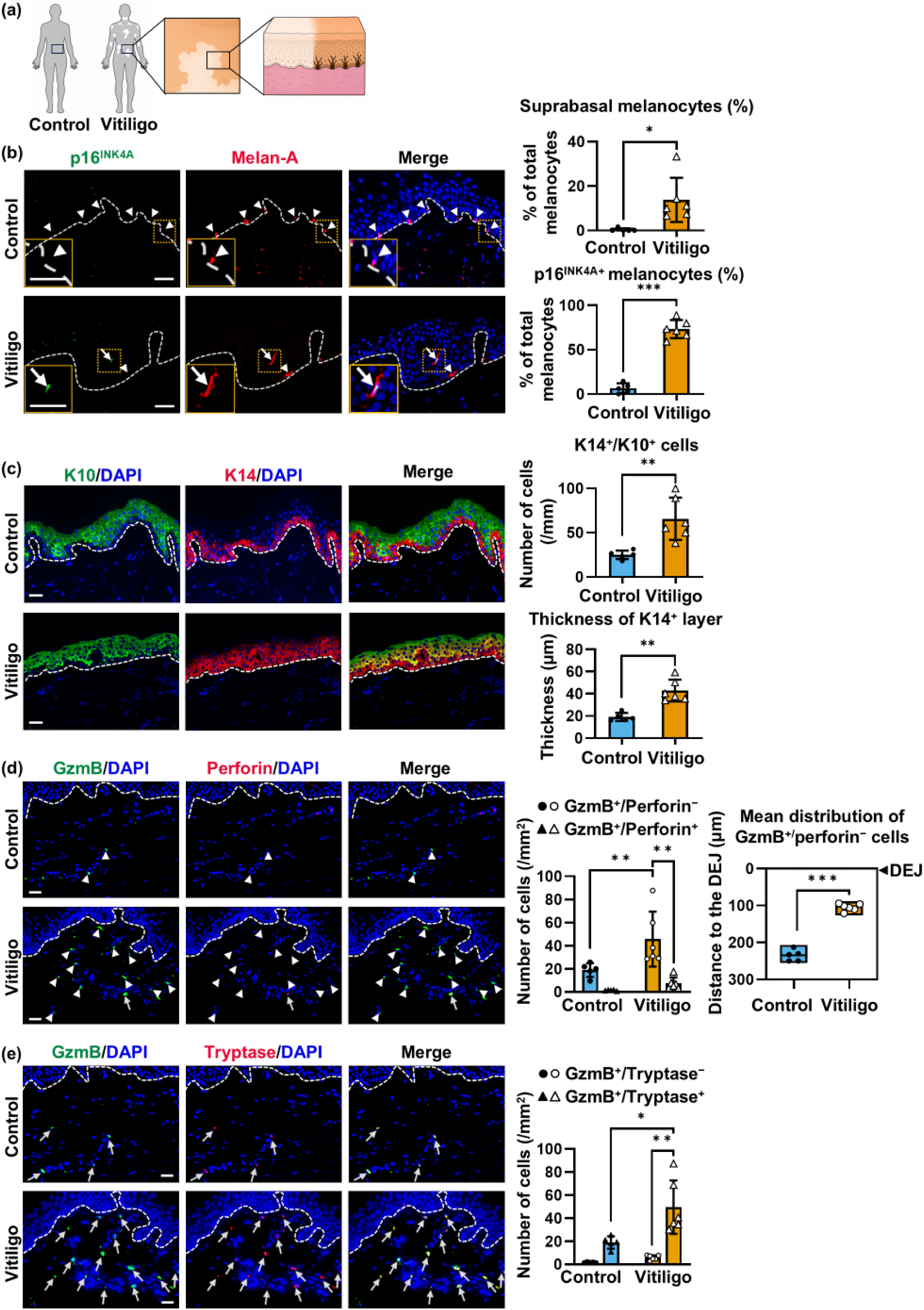
Melanocyte detachment, melanocyte senescence, aberrant keratinocyte differentiation and mast cell-associated granzyme B accumulation in human vitiligo lesion edges. (a) Skin biopsies were collected from control skin (N=5) and vitiligo skin (N=6), followed by tissue sectioning for histological analysis. (b) Representative images of vitiligo skin and control skin stained for p16^INK4A^ (green) and Melan-A (red). Arrowheads indicate basal melanocytes; arrows indicate suprabasal p16^INK4A+^ melanocytes. Melanocytes in orange boxes are enlarged in the bottom-left panels. Quantification shows the proportion of suprabasal melanocytes and p16^INK4A+^ melanocytes among total melanocytes. (c) Representative images of the vitiligo skin and control skin stained for keratin 10 (K10; green) and keratin 14 (K14; red). Quantification shows the number of K14^+^/K10^+^ cells per mm of dermal–epidermal junction (DEJ), and the mean thickness (μm) of the K14^+^ layer in the epidermis. (d) Representative images of vitiligo skin and control skin stained for granzyme B (GzmB; green) and perforin (red). White arrowheads indicate GzmB^+^/perforin^−^ cells, and arrows indicate GzmB^+^/perforin^+^ cells. Quantification shows GzmB^+^/perforin^−^ and GzmB^+^/perforin^+^ cells/mm^2^ within 300 μm of the DEJ, and the mean distance (μm) of GzmB^+^/perforin^-^ cells from the DEJ. (e) Representative images of vitiligo skin and control skin stained for GzmB (green) and tryptase (red). White arrows indicate GzmB^+^/tryptase^+^ cells. Quantification demonstrates the numbers of GzmB^+^/tryptase^−^ and GzmB^+^/tryptase^+^ cells/mm^2^ within 300 μm of the DEJ. In (b-e), 4 ,6-diamidino-2-phenylindole (DAPI) labels nuclei, and the dotted lines indicate the DEJ. Scale bars, 20 μm. Plots are shown as individual values quantified from immunofluorescence images, with mean ± SD. N = 5 for the control group and N = 6 for the vitiligo group. Statistical comparisons between two groups were performed using a two-tailed Welch’s *t*-test. \**P* ≤ 0.05, \*\**P* ≤ 0.01, \*\*\**P* ≤ 0.001 **Alt legend** Schematic of human skin biopsy collection, representative immunofluorescence images and quantitative plots comparing control and vitiligo lesional skin. Images show melanocyte localisation and p16^INK4A^ level; keratinocyte differentiation markers, keratin 10 and keratin 14; and granzyme B (GzmB) co-staining with perforin or tryptase. Vitiligo lesions show increased suprabasal and p16^INK4A^-positive melanocytes, altered keratinocyte differentiation and increased GzmB-positive cells, predominantly associated with tryptase and with limited perforin co-distribution.

**Table 1:** Clinical characteristics and immunostaining profile of vitiligo patients. Clinical information was recorded at the time of biopsy. Values represent the quantification of immunostaining images shown in Figure 1. Only cells located within 300 μm of the dermal-epidermal junction were analysed. Granzyme B (GzmB).

| Age (years) | Disease duration at biopsy | Prior therapies | Biopsy site | GzmB<sup>+</sup> cells (/mm<sup>2</sup>) | GzmB<sup>+</sup>/perforin<sup>-</sup> among GzmB<sup>+</sup> cells | GzmB<sup>+</sup>/tryptase<sup>+</sup> among GzmB<sup>+</sup> cells |
| --- | --- | --- | --- | --- | --- | --- |
| 29 | 8 months | Topical vitamin D | Clavicle | 105.2 | 83.4% | 96.8% |
| 39 | 1 month | None | Axilla | 34.5 | 89.5% | 79.3% |
| 50 | 3 years | None | Abdomen | 36.0 | 80.9% | 87.7% |
| 50 | 5 months | Topical betamethasone valerate discontinued 2 months before biopsy | Chest | 31.9 | 82.7% | 82.7% |
| 73 | 4 months | Topical betamethasone valerate | Cubital fossa | 43.3 | 87.8% | 84.0% |
| 76 | 4 months | None | Shoulder | 67.7 | 88.3% | 93.7% |

As melanocyte detachment and senescence-associated phenotype are key features of vitiligo pathology,^8–10,24^ we first assessed the localisation of the melanocyte marker Melan-A and the senescence-associated protein p16^INK4A^. Compared with control skin, active vitiligo lesions showed higher proportions of suprabasal melanocytes (14.5 vs 0.4%) and p16^INK4A^-positive melanocytes (73.3 vs 6.6%; Figure 1b).

To assess the abnormal keratinocyte differentiation and epidermal architecture alterations previously reported in vitiligo,^11,12^ we analysed keratinocyte differentiation markers, including suprabasal marker keratin 10 (K10) and basal keratinocyte marker keratin 14 (K14).^25^ In control skin, K14-positive basal and K10-positive suprabasal keratinocytes were clearly compartmentalised. However, active vitiligo lesions showed suprabasal expansion of K14-positive cells, as reflected by a thicker K14-positive layer (43.0 vs 19.1 μm) and increased K14/K10 double-positive cells (65.6 vs 25.0 cells/mm; Figure 1c). Reanalysis of a published vitiligo single-cell RNA sequencing (scRNA-seq) dataset (GSE203262) showed an expanded keratinocyte population expressing both basal keratinocyte markers *KRT5*/*KRT14* and suprabasal markers *KRT1*/*KRT10* in lesional compared with non-lesional skin (13.4 vs 2.1%; Figure S1a, b), further supporting aberrant keratinocyte differentiation in vitiligo.

We next assessed GzmB abundance and cellular distribution in vitiligo. In active lesions, double staining of GzmB and perforin showed more GzmB-positive cells in the upper dermis than control skin (53.1 vs 20.3 cells/mm^2^; Figure 1d). GzmB-positive/perforin-negative cells were closer to the dermal-epidermal junction (DEJ) (109.2 vs 234.7 μm), 85.4% of lesional GzmB-positive cells lacked perforin co-distribution (Table 1).

To define the predominant GzmB-positive population, we co-stained GzmB with cellular markers, including tryptase, CD8 and CD56 (Figure 1e, S2a-c). In active vitiligo lesions, 87.4% of GzmB-positive cells were tryptase-positive (Figure 1e), whereas only 4.4% and 3.4% were positive for CD8 and CD56, respectively (Figure S2a, b). Thus, mast cells, rather than CD8-positive T cells or natural killer cells, represented the predominant GzmB-positive population in active vitiligo lesions. Reanalysis of public scRNA-seq data (PRJCA006797) likewise showed a trend towards a higher proportion of *GZMB*-expressing/*PRF1*-null mast cells among all cells in vitiligo lesions than in control skin (0.20 vs 0.01%; Figure S2c).

These findings show that active vitiligo lesions exhibit features of melanocyte detachment, melanocyte senescence, aberrant keratinocyte differentiation and increased mast cell-associated GzmB with limited perforin association.

### Subcutaneous administration of granzyme B induces focal skin depigmentation accompanied by melanocyte detachment, melanocyte senescence and aberrant keratinocyte differentiation

To investigate whether extracellular GzmB is sufficient to induce depigmentation involving interfollicular epidermal melanocytes, we injected 100 ng active recombinant GzmB or vehicle subcutaneously into C57BL/6 mouse tails once daily (Figure 2a). By day 14, GzmB-injected skin showed focal depigmented macules and higher ΔL* values (increased skin brightness) than vehicle-injected skin (7.1 vs 1.7; Figure 2b). Whole-mount tail staining showed fewer melanocytes within keratin 31 (K31)-positive scale regions (59.6 vs 122.0 cells/scale; Figure 2c), where interfollicular melanocytes predominantly reside in mouse tail epidermis.^26,27^ To determine whether this depigmentation was associated with melanocyte detachment and senescence, we assessed vertical sections, which showed higher proportions of suprabasal melanocytes (21.5 vs 3.4%; Figure 2d) and p16^INK4A^/microphthalmia-associated transcription factor (MITF) double-positive melanocytes (71.2 vs 16.1%; Figure 2e) in GzmB-injected skin than vehicle-injected skin.

**Figure 2.**
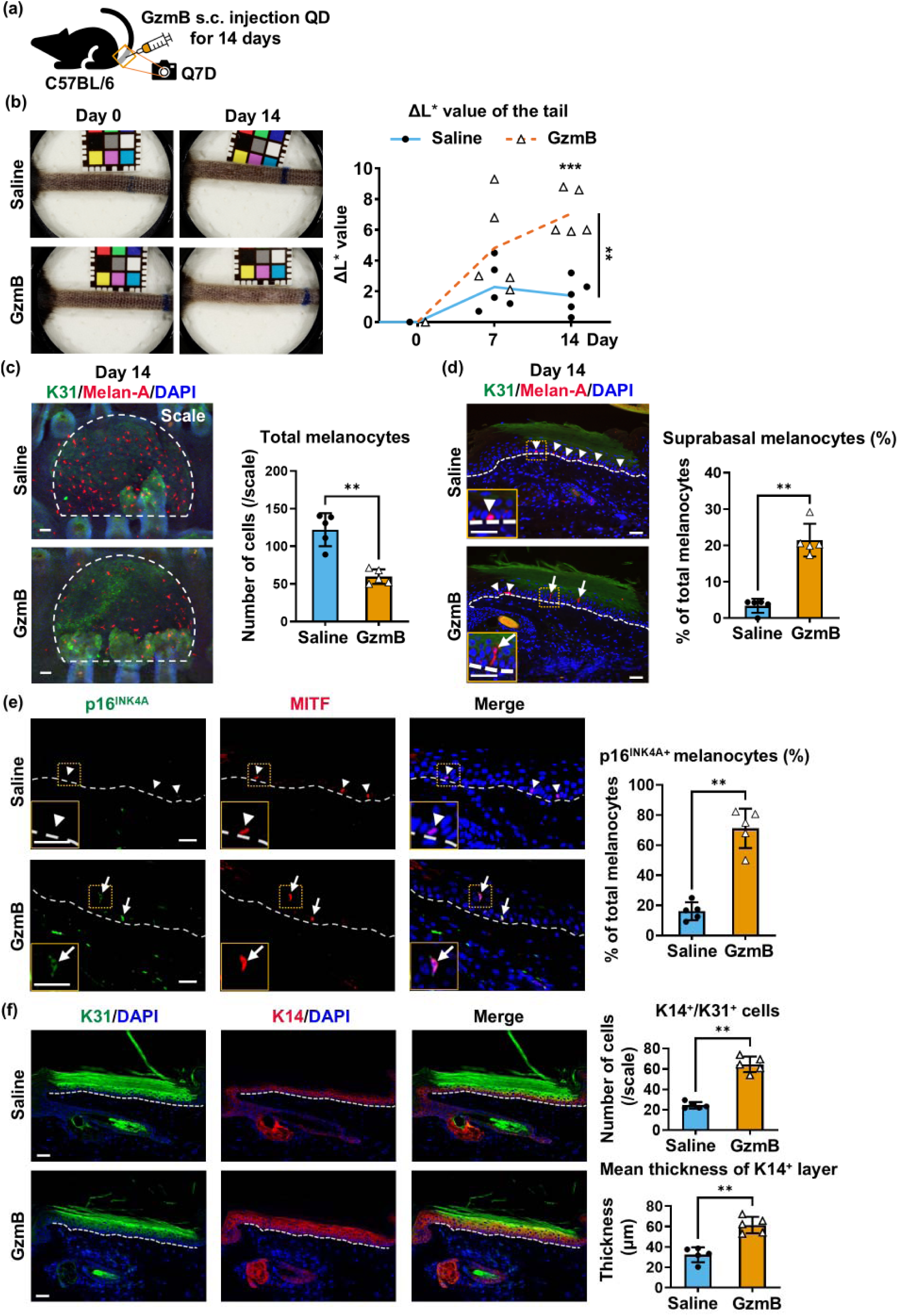
Extracellular granzyme B induces depigmentation, melanocyte detachment, melanocyte senescence and aberrant keratinocyte differentiation in mouse tail skin. (a) C57BL/6 mouse tail skin was injected subcutaneously (s.c.) with saline or granzyme B (GzmB; 100 ng) once daily (QD) for 14 days, and dermoscopic images of the tails were acquired every 7 days (Q7D). N = 5 per group. (b) Representative dermoscopic images of saline- or GzmB-injected tail skin on days 0 and 14. Quantification shows ΔL* values over time. Statistical comparisons were performed using two-way ANOVA with multiple comparisons. (c) Representative whole-mount images of saline- or GzmB-injected mouse tail epidermis stained for keratin 31 (K31; green) and Melan-A (red). White dotted lines indicate scale regions defined by K31. Quantification shows the total number of melanocytes per scale region; only melanocytes in the scale region were counted. (d) Representative vertical sections of saline- or GzmB-injected mouse tail skin stained for K31 (green) and Melan-A (red). Arrowheads indicate basal melanocytes, and arrows indicate suprabasal melanocytes. Melanocytes in orange boxes are enlarged in the bottom-left panels. Quantification shows the proportion of suprabasal melanocytes among total melanocytes within the scale regions; only melanocytes in the scale region were counted. (e) Representative images of saline- or GzmB-injected mouse tail skin stained for p16^INK4A^ (green) and microphthalmia-associated transcription factor (MITF; red). Arrowheads indicate MITF^+^/p16^INK4A−^ melanocytes, and arrows indicate MITF^+^/p16^INK4A+^ melanocytes. Melanocytes in orange boxes are enlarged in the bottom-left panels. Quantification shows the proportion of p16^INK4A+^ melanocytes among total melanocytes. (f) Representative images of saline- or GzmB-injected mouse tail skin stained for K31 (green) and keratin 14 (K14; red). Quantification shows the number of K14^+^/K31^+^ cells per scale and the mean thickness (μm) of the K14^+^ layer in the scale region. In (c–f), 4 ,6-diamidino-2-phenylindole (DAPI; blue) labels nuclei. In (d–f), dotted lines indicate the dermal–epidermal junction. Scale bars, 20 μm. In (b-f), data are shown as individual values with mean ± SD. N = 5 per group. In (c-f), statistical comparisons between two groups were performed using two-tailed Welch’s *t*-test. \**P* ≤ 0.05, \*\**P* ≤ 0.01, \*\*\**P* ≤ 0.001. **Alt legend** Schematic showing subcutaneous granzyme B (GzmB) injection into mouse tail skin. Representative dermoscopic images and quantitative plots show increased depigmentation in GzmB-injected tail skin compared with saline-injected controls. Representative whole-mount images and quantitative plots show reduced Melan-A-positive melanocyte numbers in the keratin 31 (K31)-positive scale regions; vertical immunofluorescence images and quantitative plots show higher proportions of Melan-A-positive suprabasal melanocytes and p16^INK4A^-positive melanocytes; and suprabasal expansion of keratin 14 (K14)-positive cells and increased numbers of K14/K31 double-positive cells in GzmB-injected tail skin compared with controls.

To assess whether GzmB administration also recapitulates the aberrant keratinocyte differentiation observed in vitiligo, we analysed K31, a marker of scale differentiation,^26^ together with K14. GzmB injection expanded K14-positive cells into K31-positive suprabasal layers, increased K14-positive compartment thickness (61.5 vs 32.3 μm) and K14/K31 double-positive cells (64.6 vs 24.4 cells/scale; Figure 2f) compared with vehicle-injected mice. This pattern resembled that observed in human vitiligo lesions. Survivin staining further characterised the keratinocyte differentiation pattern by assessing mitotic spindle orientation (Figure S3a). GzmB-injected skin exhibited a higher frequency of perpendicular divisions relative to the basement membrane (51.4 vs 30.5%; Figure S3b). These findings indicate increased perpendicular division, which may be associated with aberrant epidermal differentiation in vivo.

Collectively, extracellular GzmB is sufficient to induce depigmentation with vitiligo-associated pathological phenotypes, including melanocyte detachment, melanocyte senescence and abnormal keratinocyte differentiation, in mouse tail skin.

### Extracellular granzyme B reduces adhesion and induces a senescence-associated phenotype involving TGF-β/SMAD signalling in cultured melanocytes

To explore the mechanisms underlying GzmB-induced melanocyte detachment and senescence, primary neonatal human epidermal melanocytes (PHEMn) were treated with 100 nM GzmB or vehicle for 24 hours (h), with no reduction in cell viability (Figure S4a). Bulk RNA-seq identified 209 upregulated and 96 downregulated differentially expressed genes (DEGs; absolute log2 fold change (FC) > 0.2; *P* ≤ 0.05) in GzmB-treated PHEMn (Figure S5a). After redundant terms were reduced using REVIGO,^28^ Gene Ontology (GO) enrichment analysis showed that upregulated DEGs were enriched for cytokine receptor binding, whereas downregulated DEGs were enriched for chromosome segregation, mitotic cell cycle phase transition and cell-matrix adhesion (Figure 3a). Consistently, heatmaps showed downregulation of adhesion-associated DEGs and upregulation of senescence-associated secretory phenotype (SASP)-related DEGs (Figure 3b, d).

**Figure 3.**
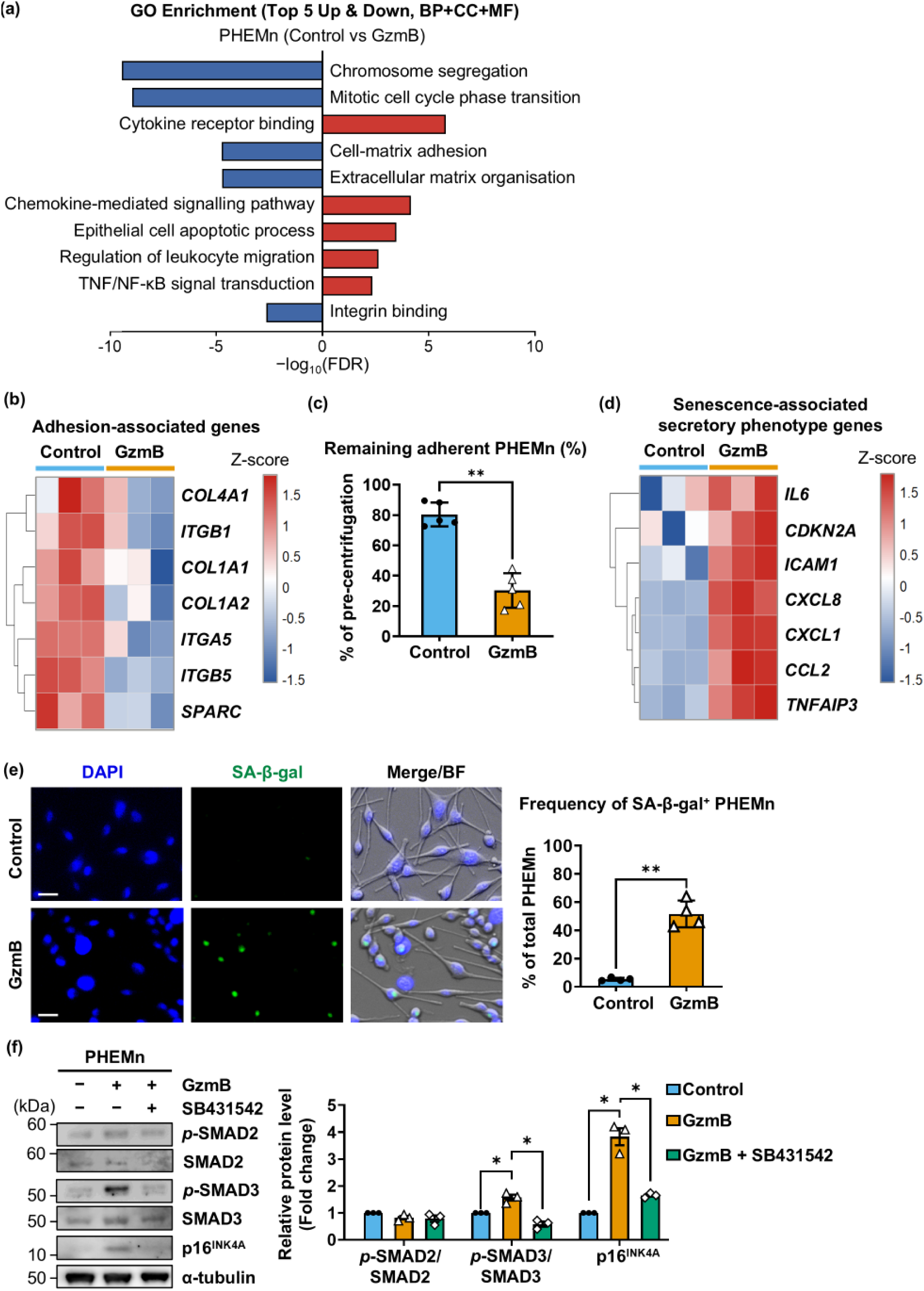
Extracellular granzyme B reduces adhesion-associated gene expression and induces a melanocyte senescence-associated phenotype involving TGF-β/SMAD signalling in vitro. (a) Gene Ontology (GO) enrichment analysis of differentially expressed genes in primary neonatal human epidermal melanocytes (PHEMn) treated with 100 nM recombinant human granzyme B (GzmB) for 24 hours (h) versus vehicle-treated controls. False discovery rate (FDR), biological process (BP), cellular component (CC), molecular function (MF). (b) Heatmap visualisation of selected adhesion-associated genes in GzmB- and vehicle-treated PHEMn. Gene expression is shown as row-scaled Z-scores. (c) Quantification of the centrifugation-based detachment assay shows the percentage of remaining cells in 24 h GzmB- and vehicle-treated PHEMn after 5 minutes of 900 *g* centrifugation. N = 5 per group. (d) Heatmap visualisation of identified senescence-associated secretory phenotype (SASP) genes in GzmB- and vehicle-treated PHEMn. Gene expression is shown as row-scaled Z-scores. (e) Representative images of PHEMn stained for senescence-associated β-galactosidase (SA-β-gal; green), 4 ,6-diamidino-2-phenylindole (DAPI; blue) and bright-field (BF). Quantification shows the frequency of SA-β-gal^+^ PHEMn. Scale bars, 10 μm. N = 4 per group. (f) Representative western blots of phosphorylated SMAD2 (*p*-SMAD2), phosphorylated SMAD3 (*p*-SMAD3), total SMAD2, total SMAD3 and p16^INK4A^ in PHEMn treated with vehicle, GzmB (100 nM) or GzmB after pretreatment with SB431542 (2 μM). Quantifications show protein levels normalised to vehicle-treated controls, which were set to 1. Statistical comparisons were performed using a two-tailed one-sample *t*-test against a hypothetical value of 1. In (a-f), N = 3 per group unless otherwise indicated. In (c, e, f), plots are shown as individual values with mean ± SD. Statistical comparisons between the two specified groups were performed using a two-tailed Welch’s *t*-test unless otherwise indicated. \**P* ≤ 0.05, \*\**P* ≤ 0.01. **Alt legend** Gene Ontology enrichment analysis and heatmap visualisation of differentially expressed genes show reduced expression of adhesion-associated genes and increased expression of senescence-associated secretory phenotype genes in granzyme B (GzmB)-treated primary neonatal human epidermal melanocytes compared with controls. Quantitative plots of the detachment assay show fewer remaining melanocytes following centrifugation after 24 hours of GzmB treatment. Representative fluorescence images and quantitative plots show increased senescence-associated β-galactosidase activity in GzmB-treated melanocytes. Immunoblot images and quantitative plots show increased SMAD2/3 phosphorylation and p16^INK4A^ protein levels following GzmB treatment, whereas SB431542 pretreatment attenuates the GzmB-induced increase in p16^INK4A^.

Given the downregulation of cell-matrix adhesion-associated DEGs, we assessed cell-matrix adhesion strength in PHEMn using a detachment assay.^29^ After 24 h of GzmB treatment followed by centrifugation (5 minutes, 900 *g*), fewer GzmB-treated than vehicle-treated cells remained adherent (29.8 vs 79.5%; Figure 3c), indicating reduced adhesion strength of GzmB-treated PHEMn.

Quantitative PCR (qPCR) also confirmed increased expression of the senescence marker *CDKN2A* and SASP genes *IL6* and *CXCL8* in GzmB-treated PHEMn (*P* ≤ 0.05; Figure S5b). An increased frequency of senescence-associated β-galactosidase (SA-β-gal)-positive cells supports a senescence-associated phenotype in GzmB-treated PHEMn (51.6 vs 5.4%; Figure 3e). Because transforming growth factor-β (TGF-β)/SMAD signalling has been implicated in cellular senescence,^30^ we examined its involvement in this phenotype. In GzmB-treated PHEMn, RNA-seq identified the TGF-β superfamily regulator *NOMO3* as the most downregulated DEG (log2FC = −1.7; *P* ≤ 0.001) and upregulated TGF/SMAD-associated DEGs including *SMAD3* (Figure S5a,c). At the protein level, activation of TGF-β/SMAD signalling was supported by a 56.7% increase in phosphorylation of SMAD3 in GzmB-treated PHEMn (Figure 3f). Mechanistically, pretreatment with TGF-β receptor inhibitor SB431542 (2 μM) attenuated GzmB-induced p16^INK4A^ upregulation (Figure 3f), supporting the involvement of TGF-β/SMAD signalling in GzmB-induced senescence-associated phenotype in PHEMn.

Together, these findings indicate that extracellular GzmB weakens adhesion and induces a senescence-associated phenotype involving TGF-β/SMAD signalling in melanocytes.

### Extracellular GzmB induces aberrant keratinocyte differentiation associated with p53 activation in vitro

To determine whether extracellular GzmB is sufficient to induce aberrant keratinocyte differentiation in vitro, primary human epidermal keratinocytes (PHEK) maintained in low-calcium conditions were treated with 100 nM GzmB for 24 h, a condition that did not reduce cell viability (Figure S4b).

RNA-seq identified 204 upregulated and 97 downregulated DEGs in GzmB-treated PHEK (Figure S5d). GO enrichment analysis demonstrated that upregulated DEGs were enriched for epidermal differentiation and regulation of cell division (Figure 4a). Consistently, the heatmap showed increased expression of differentiation-associated DEGs, including *KRT1* and *KRT10* (Figure 4b). qPCR confirmed that GzmB treatment increased *KRT1* and *KRT10* mRNA levels by 48.8 and 78.0% (Figure S5e).

**Figure 4.**
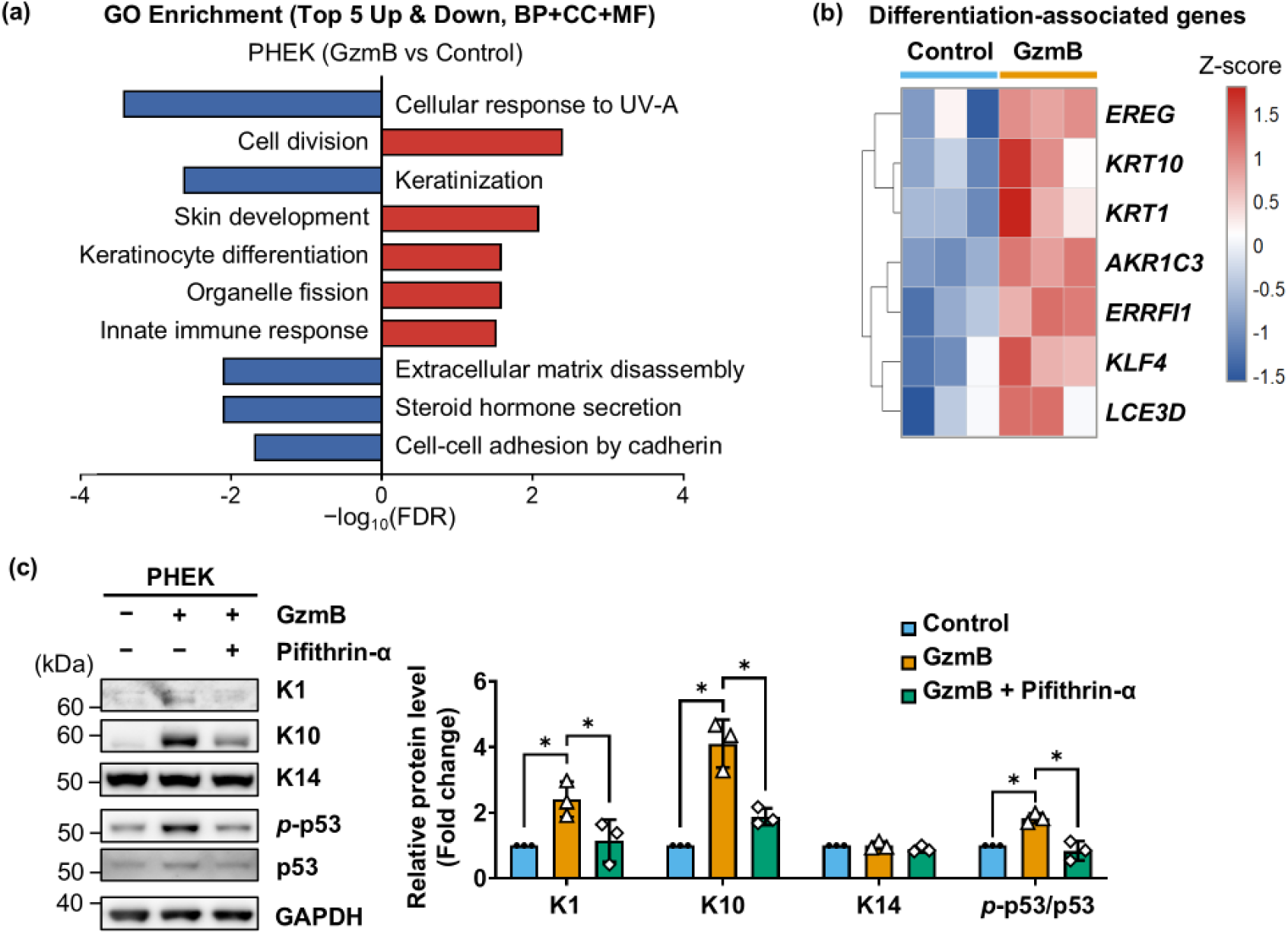
Extracellular GzmB induces aberrant keratinocyte differentiation involving p53 activation in vitro. (a) Gene Ontology (GO) enrichment analysis of differentially expressed genes in primary human epidermal keratinocytes (PHEK) treated with 100 nM recombinant human granzyme B (GzmB) for 24 hours versus vehicle-treated controls. False discovery rate (FDR), biological process (BP), cellular component (CC), molecular function (MF). (b) Heatmap visualisation of selected differentiation-associated genes in GzmB- and vehicle-treated PHEK. Gene expression is shown as row-scaled Z-scores. (c) Representative western blots of phosphorylated p53 (*p*-p53), keratin 1 (K1), keratin 10 (K10) and keratin 14 (K14) in PHEK treated with vehicle, GzmB (100 nM) or GzmB after pretreatment with pifithrin-α (5 μM). Quantifications show protein levels normalised to vehicle-treated controls, which were set to 1. Plots are shown as individual values with mean ± SD. Statistical comparisons were performed using a two-tailed one-sample *t*-test against a hypothetical value of 1. \**P* ≤ 0.05. In (a–c), N = 3 per group. **Alt legend** Gene Ontology enrichment analysis and heatmap visualisation of selected genes show increased expression of differentiation-associated and p53 signalling-associated genes in granzyme B (GzmB)-treated primary human epidermal keratinocytes compared with controls. Immunoblot images and quantitative plots show increased p53 phosphorylation and keratin 1 (K1) and keratin 10 (K10) protein levels, with retained keratin 14 (K14) levels following GzmB treatment. Pifithrin-α pretreatment attenuates GzmB-induced increases in K1 and K10 protein levels

At the protein level, keratin 1 (K1) and K10 reached 241% and 410% of control levels following GzmB treatment, whereas the basal marker K14 was maintained, supporting GzmB-induced aberrant keratinocyte differentiation (Figure 4c). Because p53 signalling is implicated in keratinocyte differentiation,^31,32^ and p53 signalling-associated DEGs were upregulated after GzmB stimulation (Figure S5f), we examined the involvement of p53 activation. Consistent with transcriptomic changes, GzmB increased p53 phosphorylation by 83.7%, while pretreatment with the p53 inhibitor pifithrin-α (5 μM) attenuated the GzmB-induced increases in K1 and K10 (Figure 4c).

These results indicate that extracellular GzmB induces aberrant keratinocyte differentiation in vitro, associated with activation of p53 signalling.

### Rhododendrol-induced leukoderma recapitulates melanocyte detachment, melanocyte senescence and aberrant keratinocyte differentiation with mast cell-associated granzyme B accumulation

To examine the functional relevance of GzmB in vitiligo-relevant depigmentation, we used an RD-induced leukoderma model. This model recapitulates depigmentation-related epidermal pathology relevant to vitiligo, but does not reproduce the autoimmune cytotoxic component of vitiligo pathology.^33^ Adapted from published protocols, 30% (w/w) RD was applied topically to mouse tails once daily for 21 days (Figure 5a).^34^

**Figure 5.**
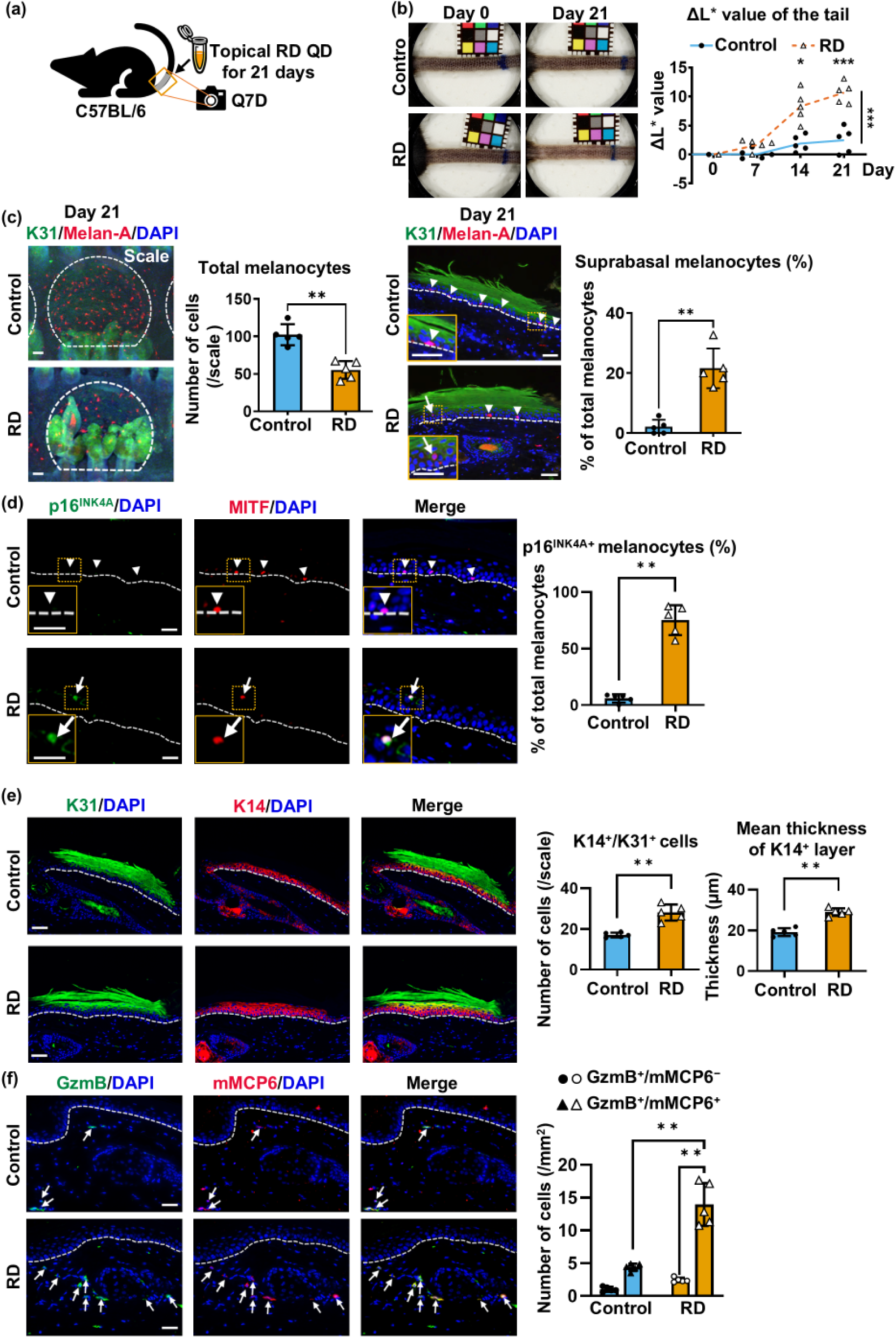
Rhododendrol-induced leukoderma recapitulates melanocyte detachment, melanocyte senescence and aberrant keratinocyte differentiation with mast cell-associated GzmB accumulation. (a) C57BL/6 mouse tail skin was treated topically with vehicle or 30% rhododendrol (RD) once daily (QD) for 21 days, and dermoscopic images of the tails were acquired every 7 days (Q7D). (b) Representative dermoscopic images of vehicle- or RD-treated mouse tail skin on days 0 and 21. Quantification shows ΔL* values over time. Statistical comparisons were performed using two-way ANOVA with multiple comparisons. (c) Representative whole-mount images of vehicle- or RD-treated mouse tail epidermis stained for keratin 31 (K31; green) and Melan-A (red). White dotted lines indicate scale regions defined by K31. Quantification shows the total number of melanocytes per scale. The right panels show representative images of vehicle- or RD-treated mouse tail skin stained for K31 (green) and Melan-A (red). Arrowheads indicate basal melanocytes, and arrows indicate suprabasal melanocytes. Quantification shows the proportion of suprabasal melanocytes among total melanocytes within the scale regions. Only melanocytes in the scale region were counted. (d) Representative images of vehicle- or RD-treated mouse tail skin stained for p16^INK4A^ (green) and microphthalmia-associated transcription factor (MITF; red). Arrowheads indicate MITF^+^/p16^INK4A−^ melanocytes, and arrows indicate MITF^+^/p16^INK4A+^ melanocytes. Melanocytes in orange boxes are enlarged in the bottom-left panels. Quantification shows the proportion of p16^INK4A+^ melanocytes among total melanocytes. (e) Representative images of vehicle- or RD-treated mouse tail skin stained for K31 (green) and keratin 14 (K14; red). Quantification shows the number of K14^+^/K31^+^ cells/scale and the mean thickness (μm) of the K14^+^ layer in the scale region. (f) Representative images of vehicle- or RD-treated mouse tail skin stained for GzmB (green) and mouse mast cell protease-6 (mMCP-6; red). White arrows indicate GzmB^+^/mMCP-6^+^ cells. Quantification shows GzmB^+^/mMCP-6^+^ cells/scale within 300 μm of the dermal–epidermal junction (DEJ). In (c–f), 4 ,6-diamidino-2-phenylindole (DAPI; blue) labels nuclei. In (d–f), dotted lines indicate the DEJ. Scale bars, 20 μm. Plots are shown as individual values with mean ± SD. N = 5 per group. Statistical comparisons between two groups were performed using a two-tailed Welch’s *t*-test. \**P* ≤ 0.05, \*\**P* ≤ 0.01, \*\*\**P* ≤ 0.001. **Alt legend** Schematic showing topical rhododendrol (RD) application to mouse tail skin. Representative dermoscopic images and quantitative plots show increased depigmentation in RD-treated tail skin compared with vehicle-treated controls. Representative whole-mount images and quantitative plots show reduced Melan-A-positive melanocyte numbers in keratin 31 (K31)-positive scale regions; vertical immunofluorescence images and quantitative plots show increased numbers of p16^INK4A^-positive melanocytes, suprabasal expansion of keratin 14 (K14)-positive cells and increased numbers of K14/K31 double-positive cells; and increased numbers of granzyme B-positive cells, most of which are associated with mouse mast cell protease-6-positive cells in RD-treated tail skin compared with controls.

RD induced tail depigmentation with increased ΔL* values compared with vehicle treatment on days 14 and 21 (8.3 vs 1.9 and 10.7 vs 2.5; Figure 5b). On day 21, whole-mount staining showed fewer melanocytes (55.3 vs 102.3 cells/scale; Figure 5c), while vertical sections demonstrated higher proportions of suprabasal melanocytes (21.6% vs 2.1%; Figure 5c) and p16^INK4A^-positive melanocytes (75.2% vs 6.0%; Figure 5d) than control skin. RD treatment also increased K14-positive layer thickness (29.0 vs 19.1 µm), K14/K31 double-positive cells (28.1 vs 17.1 cells/scale; Figure 5e), and perpendicular basal keratinocyte divisions (40.0 vs 21.0%; Figure S3c) compared with control skin. In RD-treated skin, GzmB-positive cells were increased (14.0 vs 4.4 cells/scale), with 94.0% lacking perforin co-distribution (Figure S6b), and 85.0% being mouse mast cell protease-6-positive (Figure 5f), identifying mast cells as the predominant GzmB-positive population.

Together, RD-induced leukoderma recapitulates key pathological features observed in human vitiligo lesions, including melanocyte detachment, melanocyte senescence, aberrant keratinocyte differentiation and mast cell-associated GzmB accumulation. These results support the use of this model to evaluate the role of extracellular GzmB in vitiligo-relevant depigmentation.

### Topical granzyme B inhibition attenuates rhododendrol-induced depigmentation and associated melanocyte detachment, melanocyte senescence and abnormal keratinocyte differentiation in mice

To assess the functional contribution of GzmB in vitiligo-relevant depigmentation, VTI-1002 (3.6 mg/mL), a selective GzmB inhibitor,^35^ or vehicle was applied topically to RD-treated mouse tails twice daily starting 1 day before RD application (Figure 6a).

**Figure 6.**
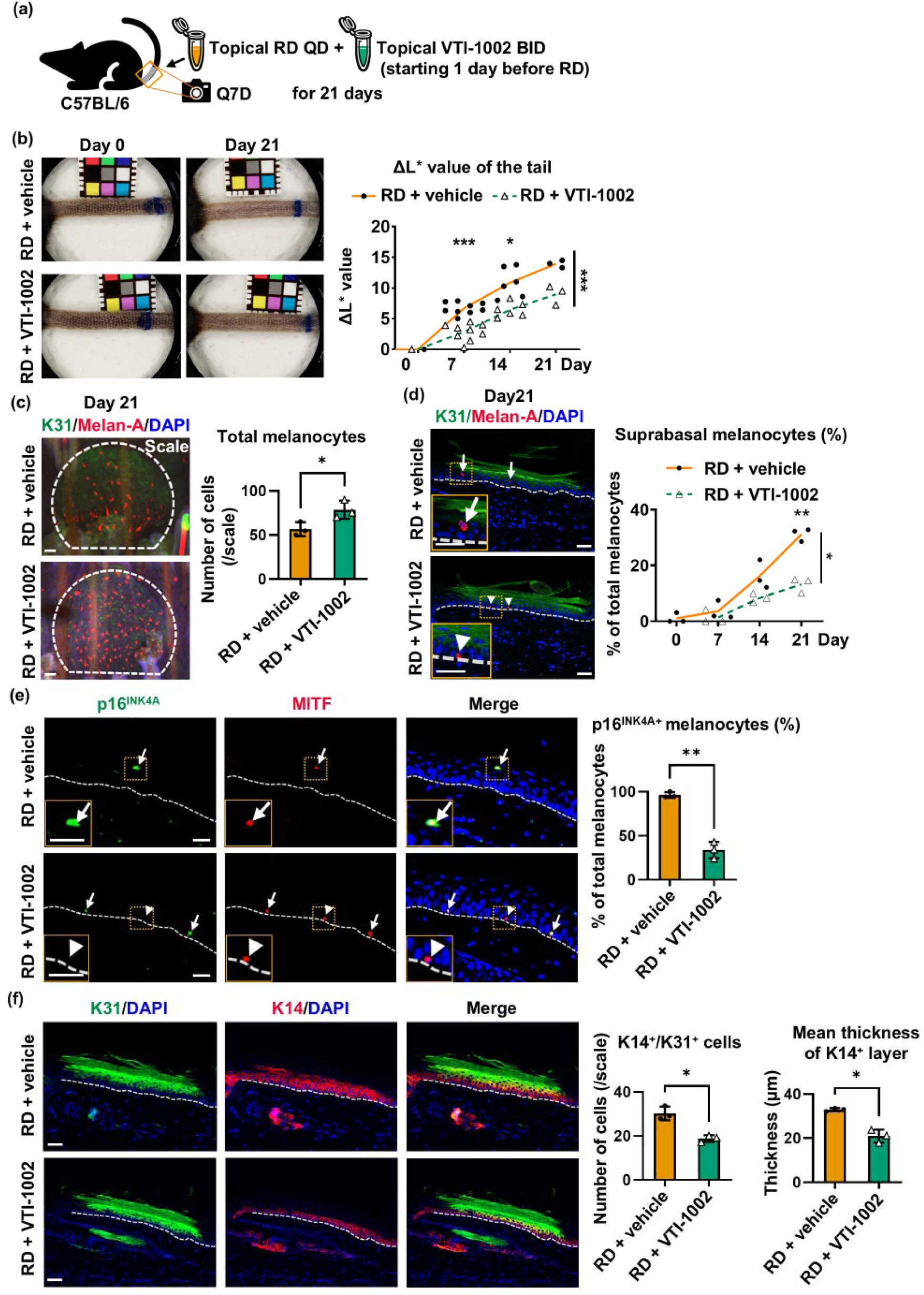
Topical GzmB inhibition attenuates rhododendrol-induced depigmentation and associated melanocyte and epidermal pathological changes. (a) In the 21-day RD-induced leukoderma model, C57BL/6 mouse tail skin was treated topically with vehicle or the granzyme B inhibitor VTI-1002 twice daily (BID), beginning 1 day before once daily (QD) rhododendrol (RD) application. Dermoscopic images were acquired every 7 days (Q7D). (b) Representative dermoscopic images of RD-treated tail skin with topical vehicle or VTI-1002 treatment on day 0 and day 21. Quantification shows ΔL* values over time. Statistical comparisons were performed using a mixed-effects two-way ANOVA with multiple comparisons. (c) Representative whole-mount images of mouse tail epidermis from the RD-induced leukoderma model treated topically with vehicle or VTI-1002, stained for keratin 31 (K31; green) and Melan-A (red). White dotted lines indicate scale regions defined by K31. Quantification shows the total number of melanocytes per scale. Only melanocytes in the scale region were counted. (d) Representative images of mouse tail skin from the RD-induced leukoderma model treated topically with vehicle or VTI-1002, stained for K31 (green) and Melan-A (red). Arrowheads indicate basal melanocytes, and arrows indicate suprabasal melanocytes. Melanocytes in orange boxes are enlarged in the bottom-left panels. Quantification shows the proportion of suprabasal melanocytes among total melanocytes within the scale regions over time. Statistical comparisons between two groups were performed using a two-way ANOVA with multiple comparisons; only melanocytes in the scale region were counted. (e) Representative images of mouse tail skin from the RD-induced leukoderma model treated topically with vehicle or VTI-1002, stained for p16^INK4A^ (green) and microphthalmia-associated transcription factor (MITF; red). Arrowheads indicate MITF^+^/p16^INK4A-^ melanocytes, and arrows indicate MITF^+^/p16^INK4A+^ melanocytes. Melanocytes in orange boxes are enlarged in the bottom-left panels. Quantification shows the proportion of p16^INK4A^-positive melanocytes among total melanocytes. (f) Representative images of mouse tail skin from the RD-induced leukoderma model treated topically with vehicle or VTI-1002, stained for K31 (green) and K14 (red). Quantification shows the number of K14^+^/K31^+^ cells per scale and the mean thickness (μm) of the K14^+^ layer in the scale region. In (c–f), 4 ,6-diamidino-2-phenylindole (DAPI; blue) labels nuclei. In (d–f), dotted lines indicate the dermal–epidermal junction. Scale bars, 20 μm. Data are shown as individual values with mean ± SD. N = 3 per group at each time point. Statistical comparisons between two groups were performed using two-tailed Welch’s *t*-test. \**P* ≤ 0.05, \*\**P* ≤ 0.01, \*\*\**P* ≤ 0.001. **Alt legend** Schematic showing topical VTI-1002 treatment in the 21-day rhododendrol (RD)-induced leukoderma mouse model. Representative dermoscopic images and quantitative plots show reduced depigmentation over time in VTI-1002-treated tail skin compared with vehicle-treated controls. Representative whole-mount images and quantitative plots show preservation of Melan-A-positive melanocytes in keratin 31 (K31)-positive scale regions. Vertical immunofluorescence images and quantitative plots show reduced proportions of Melan-A-positive suprabasal melanocytes and p16^INK4A^-positive melanocytes, together with reduced K14-positive epidermal thickening and fewer K31 and K14 double-positive cells following VTI-1002 treatment.

Topical VTI-1002 reduced RD-induced depigmentation severity with lower ΔL* values than vehicle treatment on days 7 and 14 (2.8 vs 6.7 and 6.5 vs 10.9; Figure 6b). On day 21, whole-mount staining showed that VTI-1002 preserved melanocytes compared with vehicle treatment (78.5 vs 56.7 cells/scale; Figure 6c). Vertical sections showed that the proportion of suprabasal melanocytes increased over time in both groups but remained lower in VTI-1002-treated than vehicle-treated skin on day 21 (13.2 vs 31.4%; Figure 6d). Compared with vehicle, topical VTI-1002 reduced the proportion of p16^INK4A^-positive melanocytes (33.8% vs 96.3%; Figure 6e) and reduced the thickness of the K14-positive layer (21.0 vs 33.0 μm) and K14/K31 double-positive cells (18.9 vs 30.3 cells/scale; Figure 6f). Consistently, VTI-1002-treated skin showed a lower frequency of perpendicular basal keratinocyte divisions (25.0 vs 40.0%; Figure S3d).

Collectively, pharmacological inhibition of extracellular GzmB attenuates RD-induced depigmentation and associated pathological changes, including melanocyte detachment, melanocyte senescence and aberrant keratinocyte differentiation. These features overlap with those observed in human vitiligo lesions, supporting extracellular GzmB as a functionally relevant mediator of vitiligo-associated epidermal pathology.

## Discussion

Vitiligo is a refractory and complex depigmenting disease characterised by multiple epidermal pathological features, including melanocyte detachment, melanocyte senescence and abnormal keratinocyte differentiation. Although each feature has been implicated in depigmentation, the upstream mechanisms coordinating them remain unclear. Here, we demonstrate that extracellular GzmB reproduces these pathological features in mice and induces corresponding melanocyte and keratinocyte phenotypes in vitro. GzmB was also increased and predominantly associated with perforin-negative mast cells in active human vitiligo lesions and the RD-induced leukoderma mouse model. Topical GzmB inhibition attenuated depigmentation and associated epidermal pathology in the RD model. These findings support extracellular GzmB as an upstream mediator coordinating multiple pathological features and contributing to vitiligo-associated depigmentation.

While CD8-positive T cells are key cellular mediators in vitiligo pathogenesis, accumulating evidence also implicates lesional mast cells.^36,37^ Their infiltration and activation are associated with disease activity,^38^ while elevated KIT ligand may promote mast cell recruitment.^38^ However, how mast cells contribute to melanocyte loss remains incompletely defined. Recent studies have identified mast cell-derived mediators in vitiligo, including interferon-γ (IFN-γ) and tryptase.^38^ Our findings extend this framework by identifying GzmB as an additional mast cell-associated protease in active vitiligo lesions. As mast cell-derived GzmB can act extracellularly in a perforin-independent manner,^39,40^ we examined whether it contributes to melanocyte detachment and other epidermal changes in vitiligo.

Melanocytorrhagy, defined as melanocyte detachment from the basal layer, followed by transepidermal elimination, has been implicated in melanocyte loss and depigmentation in vitiligo.^41–43^ Matrix metalloproteinase-9 (MMP-9)-mediated E-cadherin degradation promotes melanocyte detachment;^8^ however, potential additional mechanisms remain poorly explored. In the present study, extracellular GzmB downregulated cell-matrix adhesion-associated genes and decreased attachment strength in PHEMn. In the RD model, VTI-1002 reduced the proportion of suprabasal melanocytes. Together with previous MMP-9 studies, our findings support a model in which extracellular proteases, including GzmB, weaken melanocyte adhesion, thereby promoting melanocyte detachment and depigmentation in vitiligo.

Melanocyte senescence is increasingly recognised in vitiligo skin.^10,24^ Senescent melanocytes exhibit impaired melanogenic function and secrete SASP factors,^44^ contributing to both melanocyte dysfunction and a proinflammatory microenvironment.^45,46^ In the present study, p16^INK4A^-positive melanocytes were increased in vitiligo and RD-induced leukoderma lesions, and reduced by topical VTI-1002 in the RD model. In vitro, extracellular GzmB induced a senescence-associated phenotype in melanocytes, including increased *CDKN2A*, SASP-related gene expression and SA-β-gal activity. TGF-β receptor inhibition attenuated GzmB-induced p16^INK4A^ upregulation, consistent with the established cytostatic effect of TGF-β/SMAD signalling in the melanocyte lineage.^47,48^ Oxidative stress, IFN-γ, tumour necrosis factor-α and fibroblast-derived Dickkopf-1 have been implicated as upstream contributors to melanocyte senescence in vitiligo;^10,49–51^ our data identify extracellular GzmB as an additional potential trigger of melanocyte senescence.

Keratinocyte abnormalities are also indicated in vitiligo pathogenesis.^11,52^ Previous studies have identified dysregulated and stress-associated keratinocyte differentiation trajectories in vitiligo.^12,53^ Consistently, we observed increased suprabasal keratinocytes co-expressing both basal and differentiation markers in active vitiligo lesions. Extracellular GzmB reproduced this aberrant differentiation pattern in mouse epidermis and cultured PHEK, suggesting a contribution of GzmB to keratinocyte dysregulation in vitiligo. Mechanistically, GzmB induced p53-dependent aberrant keratinocyte differentiation and increased perpendicular divisions of basal keratinocytes in vivo. These findings are consistent with previous reports showing increased p53 signalling in vitiligo lesions,^54^ and a link between p53 activation and perpendicular asymmetric keratinocyte stem cell division.^55^ Given that perpendicular keratinocyte division promotes elimination of adjacent melanocytes and depigmentation in mice,^13^ the extracellular GzmB-associated perpendicular keratinocyte division may facilitate the upward displacement of neighbouring melanocytes and thereby contribute to melanocytorrhagy, particularly when their cell-matrix attachment is impaired.

Current vitiligo treatments often result in incomplete repigmentation.^56,57^ Despite the development of systemic Janus kinase inhibitors and other immunomodulators,^1^ topical therapies with limited systemic exposure are particularly desirable in this chronic, non-life-threatening disease. Topical inhibition of extracellular GzmB with VTI-1002 may therefore offer a distinct local therapy strategy. This small-molecule inhibitor selectively blocks extracellular GzmB activity, exhibits prolonged skin retention, minimal systemic exposure and no overt toxicity in mice.^35^ In the RD model, topical VTI-1002 attenuated depigmentation progression, melanocyte detachment, melanocyte senescence and abnormal epidermal differentiation. Although initiated by chemical melanocyte injury,^33^ the RD model reproduces key pathological features and mast cell accumulation observed in human vitiligo. The mast cell accumulation may reflect a compensatory increase in keratinocyte-derived SCF (KIT ligand) following melanocyte loss, consistent with other mouse models and human vitiligo.^36^ As the RD model does not recapitulate CD8-positive T cell-mediated autoimmune cytotoxicity, these findings do not diminish the importance of targeting cytotoxic immune pathways in vitiligo. Rather, topical GzmB inhibition may complement existing immunomodulatory approaches by targeting extracellular GzmB activity not addressed by current therapies.

In addition to the lack of CD8-positive T cell-mediated cytotoxicity in the RD model, several limitations warrant consideration. VTI-1002 was applied before RD induction, representing prophylactic inhibition rather than treatment of established depigmentation. The contribution of each attenuated epidermal phenotype to depigmentation was not directly assessed. The efficacy and safety of topical GzmB inhibition require validation in vitiligo patients.

Our findings expand current understanding of extracellular protease mechanisms in vitiligo-associated depigmentation. Beyond its classical intracellular cytotoxic function, GzmB promoted melanocyte detachment, melanocyte senescence and aberrant keratinocyte differentiation via an extracellular mechanism. Topical GzmB inhibition attenuated the progression of RD-induced depigmentation and associated epidermal pathology in vivo, supporting extracellular GzmB as a potential local therapeutic target for vitiligo-associated depigmentation.

## Acknowledgements

We thank Mr. Keisuke Inoue and Ms. Emi Donoue from the Research Support Platform in the Graduate School of Medicine at Osaka Metropolitan University for their assistance with sample preparation for immunohistochemistry.

## Plain Language Summary

### How granzyme B may contribute to loss of skin colour in vitiligo

Vitiligo is a skin condition in which melanocytes (pigment-producing cells) are lost, causing white patches. It affects about 0.5–2% of people worldwide. Although immune cells can attack melanocytes in vitiligo, other processes may also contribute to pigment loss.

In this study from Japan, we investigated whether granzyme B (a protein that some immune cells use to kill target cells) can promote pigment loss without directly killing melanocytes.

We analysed skin samples from patients with vitiligo and publicly available single-cell gene data. We also studied mice with pigment loss and performed laboratory experiments using human skin cells.

We found increased levels of granzyme B in vitiligo skin, mainly associated with mast cells (immune cells involved in inflammation) rather than cytotoxic T cells (immune cells that kill target cells). In laboratory experiments, granzyme B caused the detachment and premature ageing of melanocytes and disrupted the normal state of neighbouring keratinocytes (the main cells in the outer layer of skin). Applying a topical gel that blocks granzyme B to the skin of mice reduced pigment loss and associated cellular changes.

Our findings suggest that granzyme B may contribute to pigment loss through effects other than directly killing melanocytes. Blocking granzyme B outside cells could offer a new approach to treating vitiligo. Further research is needed to determine whether this approach is feasible for vitiligo patients.

