## Supporting information for "Extracellular Granzyme B Promotes Melanocytorrhagy, Melanocyte Senescence and Aberrant Epidermal Differentiation in Vitiligo and Is Therapeutically Targetable"

Materials and methods

**Human samples**

Formalin-fixed paraffin-embedded skin samples were obtained from patients with active non-segmental vitiligo lesions and from control patients at Osaka Metropolitan University Hospital (OMUH). Active vitiligo was defined as the development of new lesions or enlargement of existing lesions within the previous 6 months. Clinical information, including age, biopsy site, disease duration, disease activity and prior therapies, as shown in Table 1, was recorded at the time of biopsy. Control skin samples were collected from random skin biopsies of clinically normal-appearing sites performed to assess possible cutaneous involvement by malignant lymphoma. Histopathological examination of the biopsy showed no evidence of lymphoma, inflammatory skin disorders or depigmenting diseases (Table S1).

Written informed consent was obtained from participants who underwent skin biopsy after study initiation. For participants whose biopsy specimens had been collected before study initiation, the requirement for written informed consent was waived, and an opt-out procedure was approved by the institutional review board of OMUH. All experimental procedures involving human samples were approved by the institutional review board of OMUH (approval number: 2024-110).

**Laboratory mice**

Six-week-old female C57BL/6NCrSLC mice were purchased from Japan SLC, Inc. (Shizuoka, Japan) and housed under specific-pathogen-free conditions with a 12-hour (h) light/dark cycle, an ambient temperature of 20–26 °C, relative humidity of 40–60% and free access to food and water. After a one-week acclimation period, mice were used for the experiments described below. All experimental procedures involving animals were approved by the Osaka Metropolitan University Animal Experiment Committee (approval number: 23068).

**Murine subcutaneous granzyme B injection model**

Seven-week-old mice were anaesthetised with isoflurane (099-06571; FUJIFILM Wako Pure Chemical Corporation, Osaka, Japan), and 100 ng recombinant human active granzyme B (GzmB; GTX17622-pro; GeneTex, Inc., Irvine, CA, USA) diluted in 10 μL sterile saline (K1F91; Otsuka Pharmaceutical Factory, Inc., Tokushima, Japan) or an equal volume of saline alone was injected subcutaneously into the tail skin once daily for 14 days. Dermoscopic images were acquired on days 0, 7 and 14 using a dermoscope (DZ-D100; CASIO Computer Co., Ltd., Tokyo, Japan). Tail skin samples were collected on day 14. N = 5 per group.

**Murine rhododendrol-induced leukoderma model**

Rhododendrol (RD; R0121; Tokyo Chemical Industry Co., Ltd., Tokyo, Japan)-induced leukoderma was established with modifications from previous studies.^1,2^ Briefly, RD was prepared as a 30% (w/w) solution in 50% (v/v) ethanol (057-00451; FUJIFILM Wako). 25 μL of RD solution or vehicle alone was topically applied once daily to the proximal tail skin, covering approximately 1.5 centimetres (cm) from the tail base, for 21 days. To prevent licking of treated tail skin, mice were acclimatised to Elizabethan collars for 1 week before treatment initiation and wore the collars throughout the experiment. Dermoscopic images were acquired on days 0, 7, 14 and 21. Tail skin samples were collected on day 21 unless otherwise stated.

For GzmB inhibition experiments in the RD-induced leukoderma mouse model, VTI-1002, a GzmB-selective small-molecule inhibitor provided by viDA Therapeutics Inc. (Vancouver, BC, Canada), was formulated at 3.6 mg/mL in a gel vehicle consisting of propylene glycol (P355-1; Fisher Chemical, Thermo Fisher Scientific, Waltham, MA, USA), methylparaben (ME163; Spectrum Chemical Mfg. Corp., New Brunswick, NJ, USA) and propylparaben (PR133; Spectrum Chemical Mfg. Corp.) in 100 mM acetate buffer (pH 5.0; prepared in-house using 17.5 M glacial acetic acid [AX0073-9; EMD Millipore, Billerica, MA, USA] and sodium acetate [S1429; Sigma-Aldrich, St Louis, MO, USA]); triethanolamine (T350; Fisher Chemical) was used to neutralise the gel, as described previously.^3^ 25 μL VTI-1002 gel or vehicle gel was applied topically to mouse tail skin twice daily, beginning one day before RD treatment and continuing throughout the experiment. Dermoscopic images were obtained every 7 days. For tissue time-course analyses, tail skin samples were collected every 7 days, with N = 3 mice per group at each time point.

**Tissue preparation and immunohistochemical staining**

Mouse skin samples were fixed in 4% paraformaldehyde (163-20145; FUJIFILM Wako) and embedded in paraffin. Human and mouse paraffin-embedded samples were sectioned at 4 μm thickness, deparaffinised and rehydrated. Antigen retrieval was performed by heating sections at 95–100°C for 20 minutes (min) in citrate buffer (pH 6.1; S1699; Sigma-Aldrich). For staining protocols including nuclear antigens, sections were permeabilised with 0.1% Triton X-100 (T8787; Sigma-Aldrich) in phosphate-buffered saline (PBS; 162-19321; FUJIFILM Wako) for 10 min at room temperature; this step was omitted for protocols targeting non-nuclear antigens only. Sections were then blocked with 5% normal goat serum (NGS; 50062Z; Thermo Fisher Scientific) or 5% normal donkey serum (NDS; D9663; Sigma-Aldrich) containing 1% bovine serum albumin (BSA; 015-15103; FUJIFILM Wako) in PBS for 1 h at room temperature and incubated with primary antibodies diluted in 5% BSA in PBS overnight at 4 °C. Sections were subsequently incubated with fluorophore-conjugated secondary antibodies for 1 h at room temperature, counterstained with 4ʹ,6-diamidino-2-phenylindole (DAPI; FUJIFILM Wako, 340-07971) for 5 minutes and mounted with antifade mounting medium (Diagnostic BioSystems Inc., CA, USA, K024). Images were acquired using an APX100 fluorescence microscope (Evident Corporation, Tokyo, Japan).

Primary antibodies used for immunostaining included antibodies against GzmB (ab4059; Abcam, Cambridge, UK; 1:200), perforin (ab89821; Abcam; 1:20), tryptase (ab2378; Abcam; 1:500), CD8 (sc-7970; Santa Cruz Biotechnology, Dallas, TX, USA; 1:100), CD56 (sc-106; Santa Cruz Biotechnology; 1:100), Melan-A (sc-20032; Santa Cruz Biotechnology; 1:50; or ab210546; Abcam; 1:300), microphthalmia-associated transcription factor (MITF; AF5769; R&D Systems, Minneapolis, MN, USA; 1:200), p16^INK4A^ (ab108349; Abcam; 1:200), keratin 10 (K10; ab76318; Abcam; 1:250), keratin 14 (K14; ab197893; Abcam; 1:500), keratin 31 (K31; GP-HHA1; PROGEN Biotechnik GmbH, Heidelberg, Germany; 1:200), mouse mast cell protease-6 (mMCP-6; MAB3736; R&D Systems; 1:200) and survivin (2808; Cell Signaling Technology, Danvers, MA, USA; 1:500). Secondary antibodies used for immunostaining included Alexa Fluor 488-conjugated goat anti-rabbit IgG (A11008; Thermo Fisher Scientific, Waltham, MA, USA; 1:500), Alexa Fluor 594-conjugated goat anti-rabbit IgG (A11037; Thermo Fisher Scientific; 1:500), Alexa Fluor 594-conjugated donkey anti-rat IgG (A21209; Thermo Fisher Scientific; 1:500) and Alexa Fluor 488-conjugated goat anti-guinea pig IgG (ab150185; Abcam; 1:500).

**Whole-mount immunostaining of mouse tail epidermis**

Tail skin was incubated in 20 mM EDTA (pH 8.0; 06894-14; Nacalai Tesque, Kyoto, Japan) in PBS at 37°C for 1 h with gentle shaking to separate the epidermis from the dermis. Epidermal sheets were fixed in 4% paraformaldehyde overnight at 4°C, blocked in blocking buffer (1% BSA, 2.5% NDS, 2.5% NGS in PBS) for 1 h at room temperature, and incubated with primary antibodies against K31 (1:200) and Melan-A (ab210546; abcam) overnight at 4°C. Samples were then incubated with Alexa Fluor 488-conjugated goat anti-guinea pig IgG and Alexa Fluor 594-conjugated goat anti-rabbit IgG secondary antibodies (1:500) overnight at 4°C, counterstained with DAPI (1:1000) for 1 h at room temperature, mounted epidermal side up and imaged using FV10i confocal fluorescence microscope (Evident, Tokyo, Japan).

**Image quantification**

Images were prepared and analysed using ImageJ (version 1.54d; National Institutes of Health, Bethesda, MD, USA).

For human skin, the upper dermis was defined as the area within 300 μm below the dermal–epidermal junction (DEJ). The analysed area was calculated as the length of the DEJ multiplied by 300 μm, and cell density in this region was expressed as cells per mm², unless otherwise stated. Perforin-negative GzmB-positive cells were calculated as the percentage of GzmB-positive cells without detectable perforin signal among all GzmB-positive cells. Tryptase-positive, CD8-positive or CD56-positive cells among GzmB-positive cells were quantified using the same method. At least two sections from each individual were quantified. Suprabasal melanocytes were defined as Melan-A-positive or MITF-positive cells located above the basal layer. Suprabasal and p16^INK4A^-positive melanocytes were quantified as percentages of total melanocytes in the analysed epidermal region. Keratinocyte differentiation was assessed using K10 and K14 staining. K14-positive layer thickness was measured in at least two vertical sections per individual, and presented as the mean thickness of the whole section. K14/K10 double-positive cells per unit length of epidermis were quantified using the same method.

For mouse tail skin, total melanocyte number was expressed as the number of Melan-A-positive cells per K31-positive scale region. Suprabasal melanocytes were defined as Melan-A-positive cells located above the basal layer within K31-positive scale regions. Ten scale regions per sample were randomly selected for quantification. Keratinocyte differentiation was assessed using K14 and K31 staining. K14-positive layer thickness was measured in vertical sections at five randomly selected regions per sample. K14/K31 double-positive cells per scale region were quantified.

Mitotic spindle orientation was assessed by survivin staining according to previous studies.^4,5^ Briefly, basal keratinocytes undergoing mitosis were identified by survivin-positive mitotic structures. The angle between the mitotic spindle axis and the basement membrane was measured using ImageJ. Perpendicular divisions were defined as cell divisions with a spindle angle of 60–90° relative to the DEJ, whereas parallel divisions were defined as those with a spindle angle of 0–30° relative to the DEJ. At least 2 sections per sample were quantified.

**Dermoscopic imaging and colour analysis**

Mouse tail skin was imaged using a DZ-D100 dermoscope. Skin colour was calibrated using CASMATCH (CA010V001-1; BEAR Medic Corporation, Tokyo, Japan) according to the manufacturer's instructions. Colour was quantified in the CIE L*a*b* colour space. L* values were measured using ImageJ at 10 evenly distributed points along the treated region, extending from the tail base to the distal boundary of the treated area, and averaged for each mouse. ΔL* was calculated as the difference between the mean L* value at each time point and the corresponding baseline value on day 0.

**Cell culture and in vitro granzyme B stimulation**

Primary neonatal human epidermal melanocytes (PHEMn; PCS-200-012; ATCC, Manassas, VA, USA) were cultured in Medium 254 (M254500; Thermo Fisher Scientific) supplemented with PMA-free human melanocyte growth supplement-2 (S0165; Thermo Fisher Scientific). Primary human epidermal keratinocytes (PHEK; C-12003; PromoCell GmbH, Heidelberg, Germany) were cultured in keratinocyte growth medium (C-20011; PromoCell GmbH) supplemented with keratinocyte growth supplement (PromoCell, C-39016). Cells were maintained at 37 °C in a humidified atmosphere containing 5% CO2. Cells between passages 3 and 5 were used for experiments.

For GzmB stimulation, PHEMn and PHEK were treated with 100 nM active recombinant human GzmB or an equal volume of vehicle in culture medium for 24 h unless otherwise stated. For pharmacological inhibition of TGF-β receptor signalling in PHEMn, cells were pretreated with 2 μM SB431542 (S1067; Selleck Chemicals, Houston, TX, USA) for 2 h before GzmB stimulation.^6^ For pharmacological inhibition of p53 activity in PHEK, cells were pretreated with 5 μM pifithrin-α (166-23131; FUJIFILM Wako) for 2 h before GzmB stimulation.^7^ Inhibitors were maintained during GzmB stimulation.

**Cell viability assay**

PHEMn and PHEK were seeded in 96-well plates at 2.5 × 10^4^ cells per well and treated with 1, 10, 50 or 100 nM GzmB for 24 h. Cell viability was assessed using the cytotoxicity assay kit (CK04; Dojindo Laboratories, Kumamoto, Japan) according to the manufacturer’s instructions. Absorbance was measured using a multimode microplate reader (Varioskan LUX; Thermo Fisher Scientific). Cell viability was calculated relative to vehicle-treated controls.

**Centrifugation-based melanocyte detachment assay**

PHEMn were seeded in 96-well plates at 1.5 × 10^4^ cells per well and cultured overnight. The cells were washed once with PBS and treated with GzmB (100 nM) or vehicle for 24 h. After treatment, the wells were washed twice with PBS, and whole-well images were acquired using APX100. Each well was then completely filled with culture medium and sealed with MicroAmp Optical Adhesive Film (4311971; Thermo Fisher Scientific), ensuring that no air bubbles remained. The plate was covered with its lid, inverted and centrifuged with the cell layer facing the direction of the centrifugal force at 900 *g* for 5 min. After centrifugation, the wells were washed twice with PBS and imaged again. Remaining adherent PHEMn (%) was calculated in corresponding fields as the number of cells after centrifugation divided by the number before centrifugation in the same view. Five independent experiments were performed, with at least three replicate wells per condition in each experiment.

**Senescence-associated β-galactosidase activity assay**

Senescence-associated β-galactosidase (SA-β-gal)-positive cells were assessed in PHEMn using the cellular senescence detection kit (SG03; Dojindo Laboratories). Briefly, cells were treated with recombinant human GzmB or vehicle for 24 h, incubated with SA-β-gal detection reagent and DAPI according to the manufacturer’s protocol. Fluorescence images were acquired using an APX100 fluorescence microscope. SA-β-gal-positive cells were quantified as the mean percentage of positive cells among total DAPI-positive cells in five randomly selected fields per sample at ×20 magnification.

**RNA extraction and quantitative PCR**

Total RNA was extracted from cultured cells using the RNeasy Mini Kit (74104; QIAGEN, Hilden, Germany) according to the manufacturer’s protocol. RNA concentration and purity were determined using a NanoDrop 2000/2000c spectrophotometer (ND-2000; Thermo Fisher Scientific). Complementary DNA was synthesised using the ReverTra Ace qPCR RT Kit (FSQ-301; TOYOBO Co., Ltd., Osaka, Japan) on a PCR Thermal Cycler Dice (TP350; Takara Bio Inc., Shiga, Japan). Quantitative PCR was performed using GeneAce Probe qPCR Mix II (313-08823; NIPPON GENE CO., LTD., Tokyo, Japan) and TaqMan Gene Expression Assays (4453320; Thermo Fisher Scientific) on an Applied Biosystems Real-Time PCR System (7500 Fast; Thermo Fisher Scientific) according to the manufacturer’s protocol. Relative gene expression was normalised to GAPDH and expressed as fold change relative to vehicle-treated controls using the 2−ΔΔCt method. TaqMan assay IDs were as follows: IL6, Hs00174131_m1; CXCL8, Hs00174103_m1; CDKN2A, Hs00923894_m1; KRT1, Hs00196158_m1; KRT10, Hs00166289_m1; GAPDH, Hs99999905_m1.

**Bulk RNA sequencing and transcriptomic analysis**

RNA integrity was assessed using the RNA Nano 6000 Assay Kit on a 2100 Bioanalyzer (Agilent Technologies, Santa Clara, CA, USA). Library preparation, sequencing and primary bioinformatic processing were performed by Novogene Co., Ltd. (China Sequencing Center, Beijing, China). Briefly, indexed libraries were sequenced on a NovaSeq platform (Illumina, San Diego, CA, USA) to generate 150-bp paired-end reads. Reads were filtered using fastp (HaploX Biotechnology, Shenzhen, China), aligned using HISAT2 (version 2.0.5) and quantified using featureCounts (version 1.5.0-p3). Differential expression was analysed using DESeq2 (version 1.20.0) with Benjamini–Hochberg correction. Three biological replicates were analysed per group. Gene Ontology (GO) enrichment analysis and visualisation of volcano plots and heatmaps were performed in-house using R (version 4.0.0; R Foundation for Statistical Computing, Vienna, Austria). Differentially expressed genes were defined using an FDR-adjusted P ≤ 0.05 and an absolute log2 fold change > 0.2, unless otherwise specified. GO enrichment analysis was performed using clusterProfiler (version 4.12.6) based on the GO database (release 2026-06-15), and redundant terms were reduced using REVIGO (version 1.8.2; Ruđer Bošković Institute, Zagreb, Croatia).^8^ Heatmaps display row-scaled Z-scores of normalised gene expression. Adhesion-associated genes shown in heatmaps were selected from differentially expressed genes annotated to cell adhesion-, cell–matrix adhesion-, extracellular matrix organisation-, focal adhesion-, or basement membrane-related GO terms, with relevance to melanocyte attachment assessed based on published literature.^9^ Senescence-associated secretory phenotype (SASP)-related, TGF-β/SMAD signalling, keratinocyte differentiation-associated and p53 signalling related genes were identified by overlapping differentially expressed genes with published gene signatures in the molecular signatures database (MSigDB, version 2026.1.Hs).^10^

**SDS-PAGE and immunoblot analyses**

Cells were lysed in RIPA buffer (16488-34; Nacalai Tesque) containing protease inhibitor (03969-21; Nacalai Tesque) and phosphatase inhibitors (07575-51; Nacalai Tesque). Protein concentrations were determined with the BCA Protein Assay Kit (23225; Thermo Fisher Scientific), according to the manufacturer’s protocol. Equal amounts of proteins from each sample were mixed 1:1 with freshly prepared Laemmli sample buffer containing 125 mM Tris–HCl (pH 6.8; 010-17451; FUJIFILM Wako), 4% (w/v) sodium dodecyl sulfate (191-07145; FUJIFILM Wako), 20% (v/v) glycerol (075-00616; FUJIFILM Wako), 0.002% (v/v) bromophenol blue (021-02911; FUJIFILM Wako) and 10% (v/v) 2-mercaptoethanol (137-06862; FUJIFILM Wako), heated at 95 °C for 5 min, separated by sodium dodecyl sulfate-polyacrylamide gel electrophoresis (SDS-PAGE; Thermo Fisher Scientific), and transferred to polyvinylidene difluoride membranes (637-52261; FUJIFILM Wako). Membranes were blocked with 3% skimmed milk (190-12865; FUJIFILM Wako) in Tris-buffered saline (TBS; T1941; Takara Bio Inc.) supplemented with 0.05% Tween-20 (103168; MP Biomedicals, Solon, OH, USA) (TBST) and then probed with primary antibodies overnight at 4 °C. After washing with TBST, membranes were incubated with horseradish peroxidase (HRP)-conjugated secondary antibodies for 1 h at room temperature. Signals were detected using SuperSignal™ West Atto Ultimate Sensitivity Substrate (A38555; Thermo Fisher Scientific) and imaged using a luminescent image analyser (ImageQuant LAS 4000; GE Healthcare, Uppsala, Sweden).

Primary antibodies used for immunoblotting included antibodies against phosphorylated SMAD2/3 (MAB8935; R&D Systems; 1:200), total SMAD2/3 (AF3797; R&D Systems; 1:500), p16^INK4A^ (ab108349; Abcam; 1:200), phosphorylated p53 (9284; Cell Signaling Technology; 1:500), total p53 (2527; Cell Signaling Technology; 1:200), K1 (GP-K1; PROGEN Biotechnik GmbH; 1:500), K10 (ab76318; Abcam; 1:5000), K14 (ab197893; Abcam; 1:20000), α-tubulin (11224-1-AP; Proteintech Group, Rosemont, IL, USA; 1:20000) and GAPDH (2118; Cell Signaling Technology; 1:20000). Horseradish peroxidase-conjugated secondary antibodies used for immunoblotting included goat anti-rabbit IgG (ab6721; Abcam; 1:2000), goat anti-mouse IgG (1031-05; SouthernBiotech, Birmingham, AL, USA; 1:5000), goat anti-guinea pig IgG (ab6908; Abcam; 1:2000) and donkey anti-goat IgG (sc-2020; Santa Cruz Biotechnology; 1:5000). Band intensities were quantified using ImageJ. Phosphorylated proteins were normalised to the corresponding total proteins where applicable. Other proteins were normalised to α-tubulin or GAPDH. Values were then normalised to vehicle controls, which were set to 1.

**Single-cell RNA-sequencing reanalysis**

Published single-cell RNA-sequencing datasets were reanalysed to examine keratinocyte and immune-cell phenotypes. Data were analysed using Seurat (version 4.0).^11^ Cells were retained for analysis if they met the following quality-control criteria: nFeature_RNA ≤ 5000, nCount_RNA ≤ 25000 and percent.mt ≤ 10%. Data were normalised, scaled and subjected to principal component analysis (PCA), clustering and uniform manifold approximation and projection (UMAP) visualisation. Cell populations were annotated using canonical marker genes according to previous studies.^12-14^

Dataset GSE203262 was used for keratinocyte analysis and included lesional and contralateral non-lesional skin from patients with vitiligo. Keratinocyte populations co-expressing basal markers *KRT5*/*KRT14* and suprabasal differentiation markers *KRT1*/*KRT10* were quantified. As *KRT1* and *KRT10* expression showed an apparent bimodal distribution, cells with normalised expression values above the 75th percentile were defined as high-expressing cells.

Dataset PRJCA006797 was used for immune cell analysis and included skin samples from patients with vitiligo and from controls. Mast cells were identified using canonical mast-cell markers, including *KIT*, *TPSAB1*, *TPSAB2* and *CPA3*. Cells with at least one unique molecular identifier (UMI) mapped to *GZMB* were classified as *GZMB*-expressing, whereas cells with no detected *PRF1* UMIs were classified as *PRF1*-undetected.

**Statistical analysis**

Statistical analyses were performed using GraphPad Prism (version 9.5.1; San Diego, CA, USA) and R. Data are presented as individual values with mean ± standard deviation (SD) unless otherwise stated. Comparisons between two independent groups were performed using two-tailed Welch's *t*-test. Time-course ΔL* values were analysed using two-way ANOVA with multiple comparisons. For the GzmB inhibition experiment, in which animals were sacrificed at predefined time points, time-course data were analysed using a mixed-effects two-way ANOVA with treatment and time as fixed effects, as missing values arose from the planned experimental design. For immunoblot quantification, comparisons with vehicle control values normalised to 1 were performed using a two-tailed one-sample *t*-test against a hypothetical value of 1, whereas comparisons between GzmB and GzmB plus inhibitor groups were performed using a two-tailed Welch's *t*-test. P ≤ 0.05 was considered statistically significant.

Table

Table S1: Clinical profile of human controls

| **Age**  **(years)** | **Biopsy site** | **Clinical context** | **Histopathology** | **Prior therapies** |
| --- | --- | --- | --- | --- |
| 31 | Abdomen | Fever of unknown origin; lymphoma subsequently excluded. | No tumour involvement | None |
| 46 | Abdomen | Occipital white-matter lesion; Malignant lymphoma. | No tumour involvement | None |
| 70 | Abdomen | Primary central nervous system lymphoma | No tumour involvement | Flutoprazepam;  Sodium valproate |
| 70 | Abdomen | Primary central nervous system lymphoma | No tumour involvement | Vonoprazan; olmesartan medoxomil |
| 70 | Abdomen | Prostate cancer; fever of unknown origin; lymphoma subsequently excluded. | No tumour involvement | Bicalutamide;  Vibegron; rosuvastatin |

Control skin samples were obtained from random skin biopsies performed to assess possible cutaneous involvement by malignant lymphoma. Histopathological examination of the biopsy showed no evidence of lymphoma involvement, inflammatory skin disorders or depigmenting diseases. Prior therapies refer to systemic or skin-directed treatments administered before biopsy.

Figure Legend

Supplementary Figure 1: Single-cell RNA-sequencing reanalysis of keratinocyte differentiation marker expression in vitiligo lesions.

(a) Feature plots showing expression of *KRT5*, *KRT14*, *KRT1* and *KRT10* in published single-cell RNA-sequencing (scRNA-seq) data from non-lesional and lesional vitiligo skin from 6 vitiligo patients. Data were normalised, scaled and subjected to principal component analysis (PCA), clustering and uniform manifold approximation and projection (UMAP) visualisation. Dotted circles indicate keratinocyte populations with increased co-expression of basal and suprabasal markers. (b) Quantification of *KRT5*^high^/*KRT14*^high^ cells among *KRT1*^high^/*KRT10*^high^ keratinocyte populations in non-lesional and lesional vitiligo skin. Data are shown as individual values with mean ± SD. N = 6 per group. Statistical comparison between two groups was performed using two-tailed paired *t*-tests. *P ≤ 0.05.

**Alt legend**

UMAP plots, feature plots and quantitative analyses from single-cell RNA-sequencing reanalysis of keratinocytes from vitiligo lesional and contralateral non-lesional skin. Vitiligo lesions show an increased keratinocyte population co-expressing basal *KRT5* and *KRT14* and suprabasal differentiation markers *KRT1* and *KRT10*.

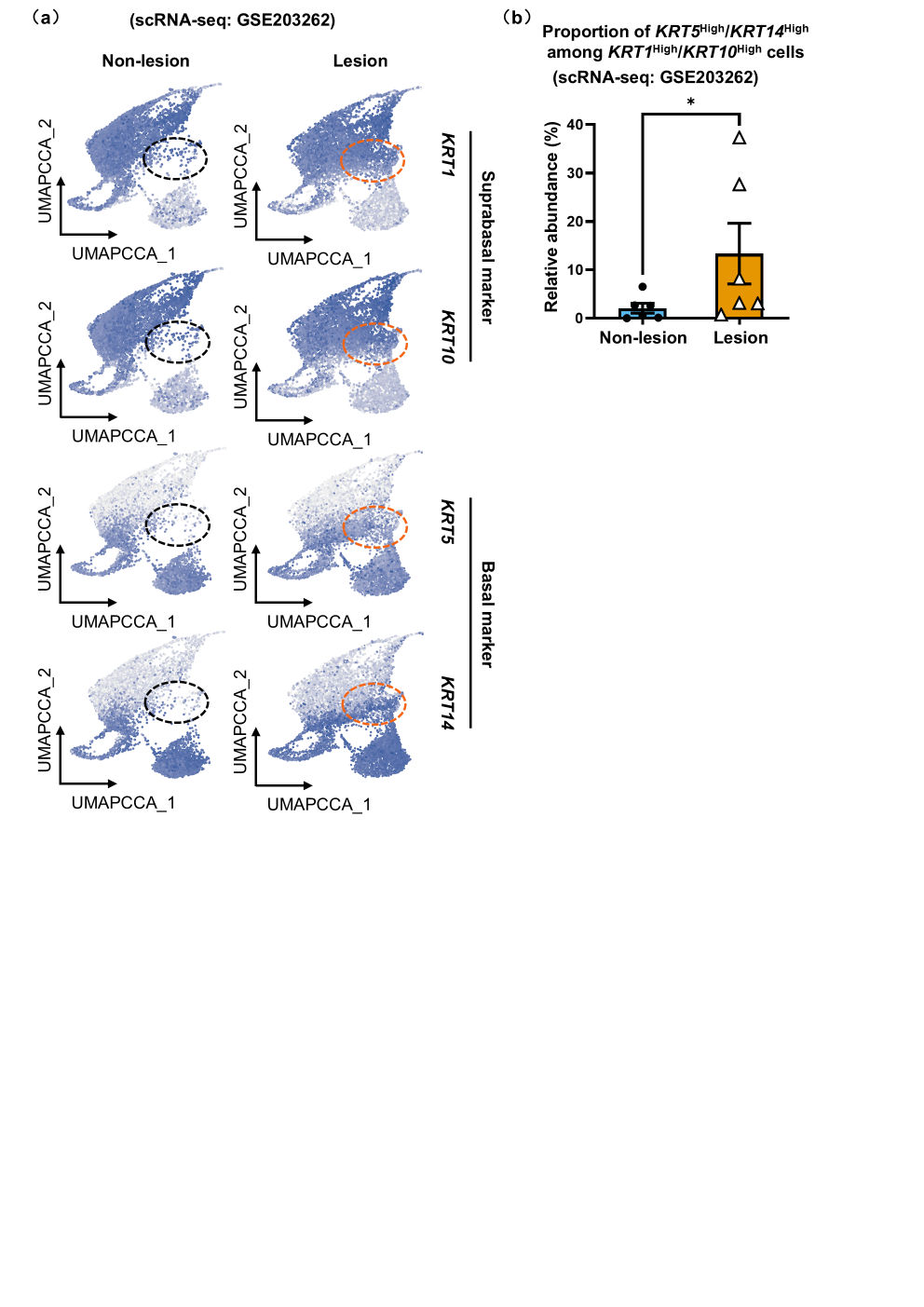

Supplementary Figure 2: GzmB shows limited co-distribution with CD8-positive and CD56-positive cells in vitiligo lesions.

(a) Representative images of human vitiligo lesional skin and control skin stained for granzyme B (GzmB; green), CD8 (red) and 4ʹ,6-diamidino-2-phenylindole (DAPI; blue). Dotted lines indicate the dermal–epidermal junction (DEJ). Scale bar, 20 µm. White arrows indicate GzmB^+^/CD8^+^ cells. Quantification shows GzmB^+^/CD8^−^ and GzmB^+^/CD8^+^ cell numbers within 300 µm of the DEJ. N = 5 for the control group and N = 6 for the vitiligo group. (b) Representative images of human vitiligo lesional skin and control skin stained for GzmB (green), CD56 (red) and 4ʹ,6-diamidino-2-phenylindole (DAPI; blue). Dotted lines indicate the DEJ. Scale bar, 20 µm. White arrows indicate GzmB^+^/CD56^+^ cells. Quantification shows GzmB^+^/CD56^−^ and GzmB^+^/CD56^+^ cell numbers within 300 µm of the DEJ. N = 5 for the control group and N = 6 for the vitiligo group. (c) Quantification of *GZMB*^+^/*PRF1*^−^ mast cells among total cell count in each individual from public single-cell RNA-sequencing (scRNA-seq) data from healthy control and vitiligo skin. N = 5 for control, N = 10 for vitiligo. (a-c) Plots show individual values with mean ± SD. Statistical comparisons between two groups were performed using two-tailed Welch’s *t*-test. **P ≤ 0.01, ***P ≤ 0.001.

**Alt legend**

Representative immunofluorescence images and quantitative plots of control and vitiligo lesional skin stained for granzyme B (GzmB) with CD8 or CD56. Most GzmB-positive cells show limited co-distribution with CD8-positive cells or CD56-positive cells. Quantification of public single-cell RNA-sequencing data from healthy control and vitiligo skin shows that mast cells with high *GZMB* and negative *PRF1* expression tended to be more abundant in vitiligo.

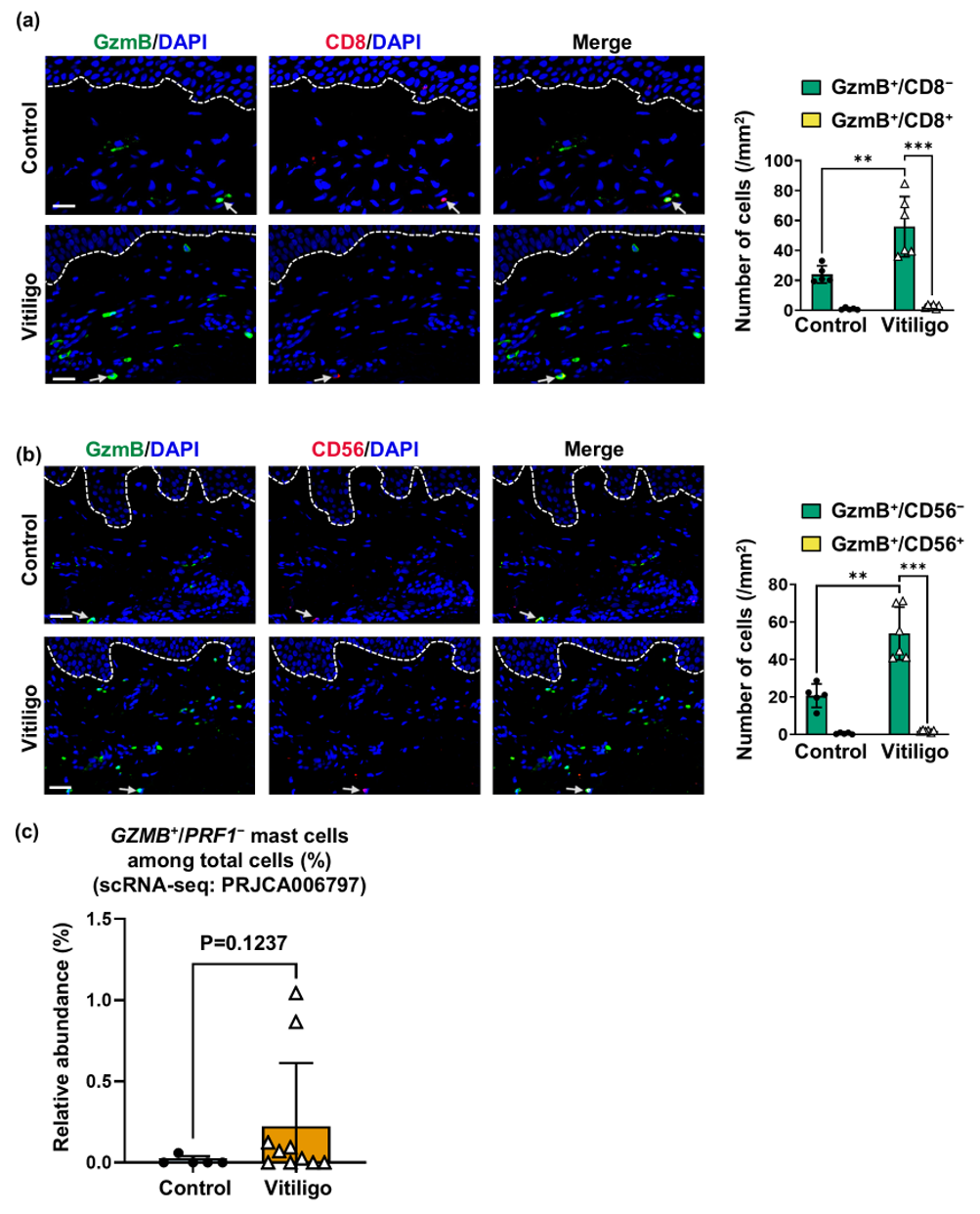

**Supplementary Figure 3:** Perpendicular division of basal keratinocytes is increased in the GzmB-injected and RD-induced depigmentation models and attenuated by VTI-1002.

(a) Representative immunostaining images for survivin (green) and 4ʹ,6-diamidino-2-phenylindole (DAPI; blue) with schematic diagrams showing perpendicular and parallel division patterns of basal keratinocytes in mouse tail epidermis. Yellow arrows indicate the angle measured between the mitotic spindle axis (yellow line) and the dermal–epidermal junction (DEJ; yellow dotted line). Scale bars, 2 μm. Perpendicular divisions were defined as cell divisions with a spindle angle of 60–90° relative to the basement membrane, whereas parallel divisions were defined as those with a spindle angle of 0–30° relative to the basement membrane. Oblique: 30-59°. (b-d) Left panels show quantified radial histograms of epidermal basal keratinocyte division angles in control mice compared with granzyme B (GzmB)-injected mice (b; N = 5) or rhododendrol (RD)-induced leukoderma mice (c; N = 5), and RD-treated mice with or without VTI-1002 (d; N = 3). The right panels show proportions of cell division types classified based on spindle angles described in (a); data are shown as mean + SD. Statistical comparisons between two groups were performed using a two-tailed Welch’s *t*-test. *P ≤ 0.05, **P ≤ 0.01.

**Alt legend**

Representative survivin immunofluorescence images, schematic diagrams, radial histograms and quantitative plots showing basal keratinocyte division orientation in mouse tail epidermis. Perpendicular divisions were more frequent in granzyme B (GzmB)-injected and rhododendrol (RD)-treated skin than in their respective controls. In the RD model, topical VTI-1002 reduced the frequency of perpendicular divisions compared with vehicle treatment.

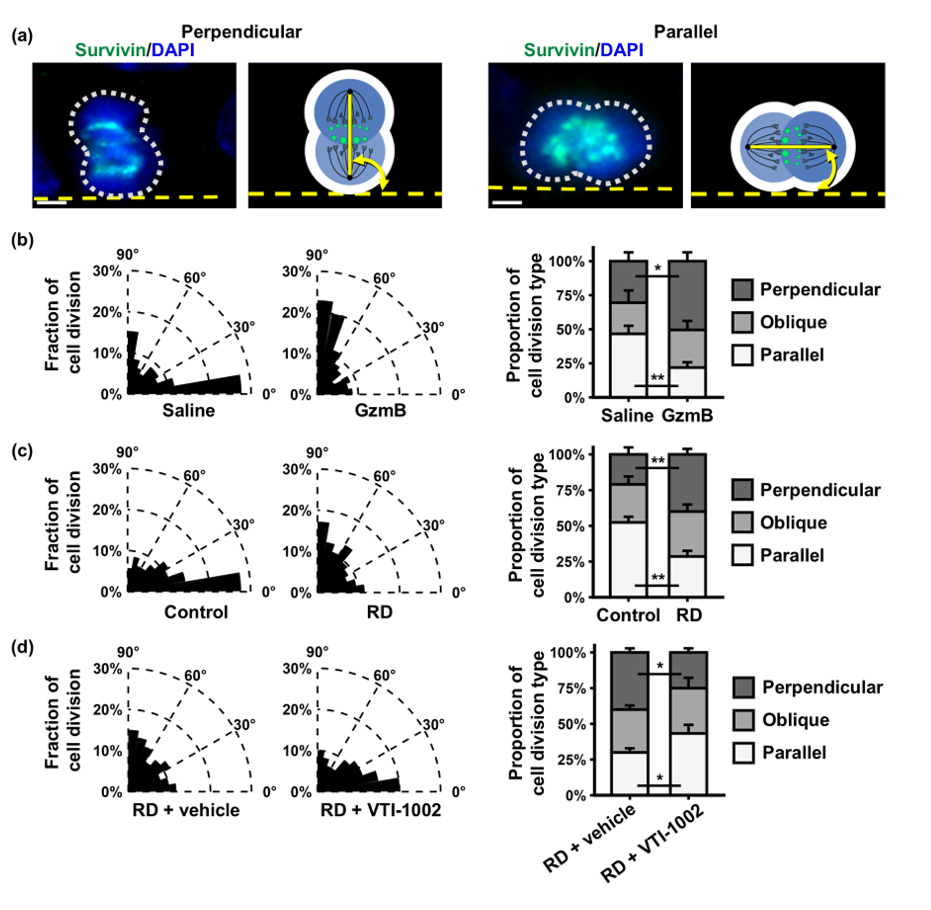

**Supplementary Figure 4:** Extracellular GzmB shows a non-cytotoxic effect on human melanocytes and keratinocytes at concentrations up to 100 nM.

(a-b) Quantification of cell viability of primary neonatal human epidermal melanocytes (PHEMn) or primary human epidermal keratinocytes (PHEK) treated with increasing concentrations of recombinant human granzyme B (GzmB) for 24 h. Data are shown as individual values with mean ± SD. N = 3 per group. Statistical comparisons were performed using one-way ANOVA followed by Dunnett’s multiple-comparisons test with 0 nM as the control. Not significant (ns), P > 0.05.

**Alt legend**

Quantitative cell viability analyses of human melanocytes and keratinocytes treated with increasing concentrations of recombinant human granzyme B (GzmB) for 24 hours. GzmB treatment at the concentration of 100 nM used for subsequent experiments does not significantly reduce cell viability.
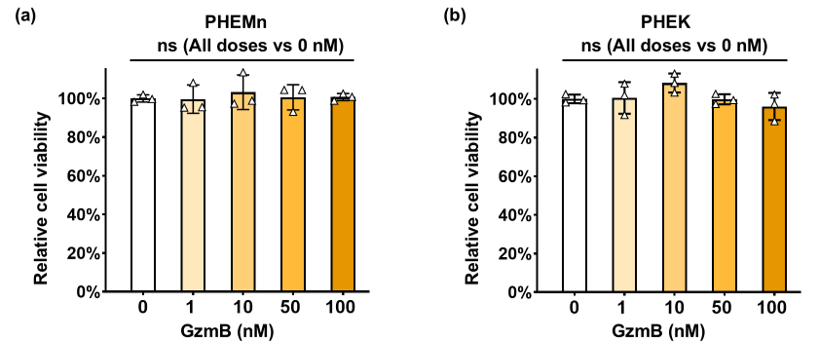

Supplementary Figure 5: Extracellular GzmB induces transcriptomic changes associated with melanocyte senescence and keratinocyte differentiation in vitro.

(a) Volcano plot showing differentially expressed genes (DEGs) in primary neonatal human epidermal melanocytes (PHEMn) treated with recombinant human granzyme B (GzmB) compared with vehicle for 24 h. False Discovery Rate (FDR), fold change (FC). N = 3. (b) Quantitative PCR (qPCR) analysis of senescence-associated genes, including *IL6*, *CXCL8* and *CDKN2A,* in control and GzmB-treated PHEMn, shown as individual values with mean ± SD. N = 5. (c) Heatmap visualisation of transforming growth factor-β (TGF-β)/SMAD-related DEGs in control and GzmB-treated PHEMn. N = 3. (d) Volcano plot showing DEGs in primary human epidermal keratinocytes (PHEK) treated with recombinant human GzmB compared with vehicle for 24 h. N = 3. (e) qPCR analysis of keratinocyte differentiation genes, including *KRT1* and *KRT10,* in vehicle- and GzmB-treated PHEK shown as individual values with mean ± SD. N = 5.

(f) Heatmap visualisation of p53 signalling-associated DEGs in control and GzmB-treated PHEK. N = 3. Statistical comparisons between two groups were performed using two-tailed Welch’s *t*-test. *P ≤ 0.05, **P ≤ 0.01.

**Alt legend**

Volcano plots show differentially expressed genes in granzyme B (GzmB)-treated primary neonatal human epidermal melanocytes and primary human epidermal keratinocytes compared with controls. Quantitative PCR analyses and heatmaps visualisation of selected differentially expressed genes show increased expression of senescence-associated genes and changes in transforming growth factor-β/SMAD-related genes in GzmB-treated melanocytes, and increased expression of differentiation-associated genes and changes in p53 signalling-associated genes in GzmB-treated keratinocytes compared with controls.

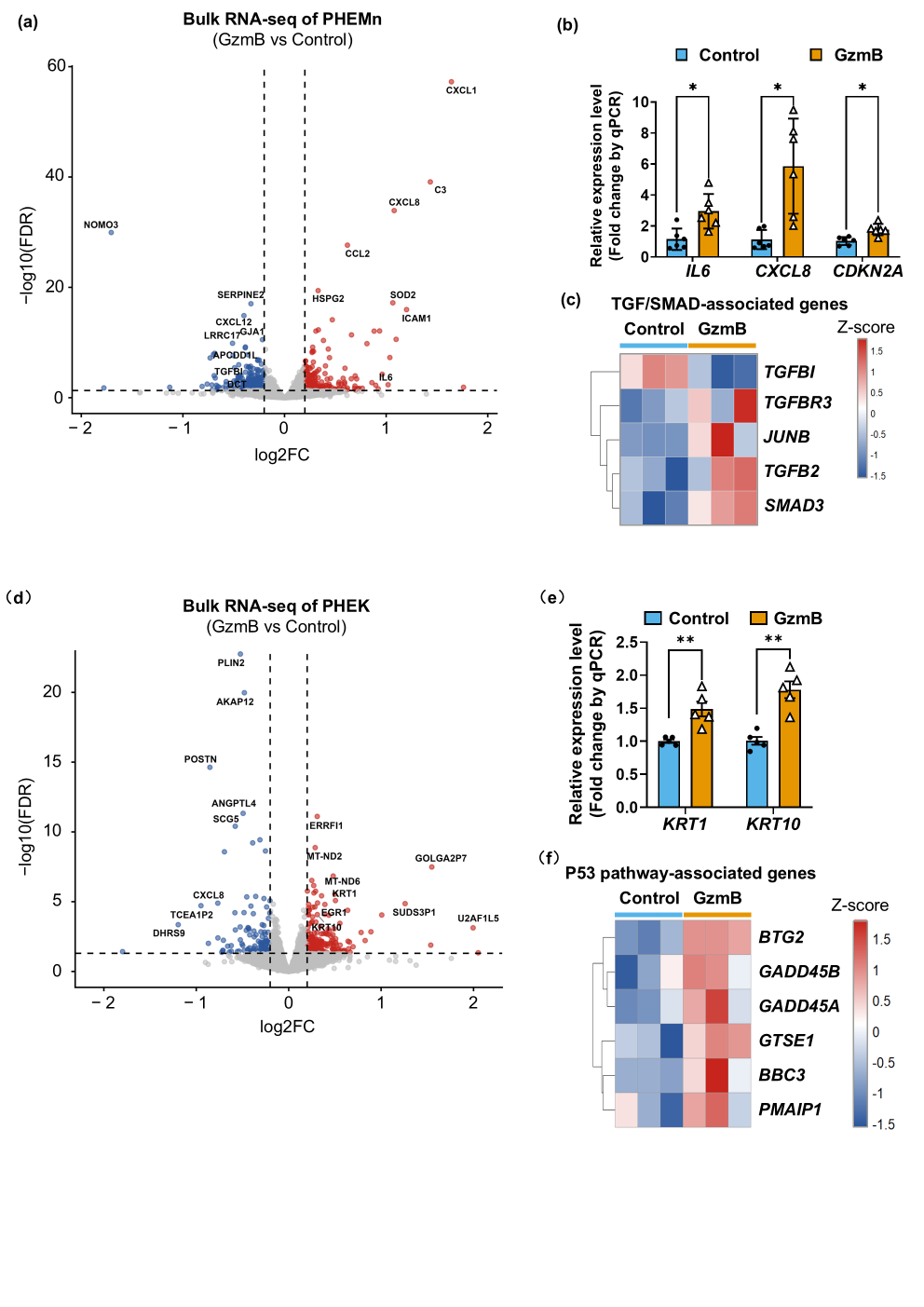

Supplementary Figure 6: RD-induced leukoderma shows increased GzmB-positive cells with limited perforin association.

(a) Representative images of mouse tail skin stained for GzmB (green), perforin (red) and 4ʹ,6-diamidino-2-phenylindole (DAPI; blue). White arrowheads indicate GzmB^+^/perforin^−^ cells. White dotted lines indicate the dermal–epidermal junction (DEJ). Quantification shows GzmB^+^/perforin^−^ and GzmB^+^/perforin^+^ cell numbers within 300 μm of the dermal–epidermal junction (DEJ). Scale bars, 40 μm. Plots are shown as individual values with mean ± SD. N = 5 per group. Statistical comparisons between two groups were performed using two-tailed Welch’s *t*-test. **P ≤ 0.01.

**Alt legend**

Representative immunofluorescence images and quantitative plots of mouse tail skin during rhododendrol-induced leukoderma. Rhododendrol treatment increased the numbers of granzyme B (GzmB)-positive cells, with most GzmB-positive cells showing limited perforin co-localisation.
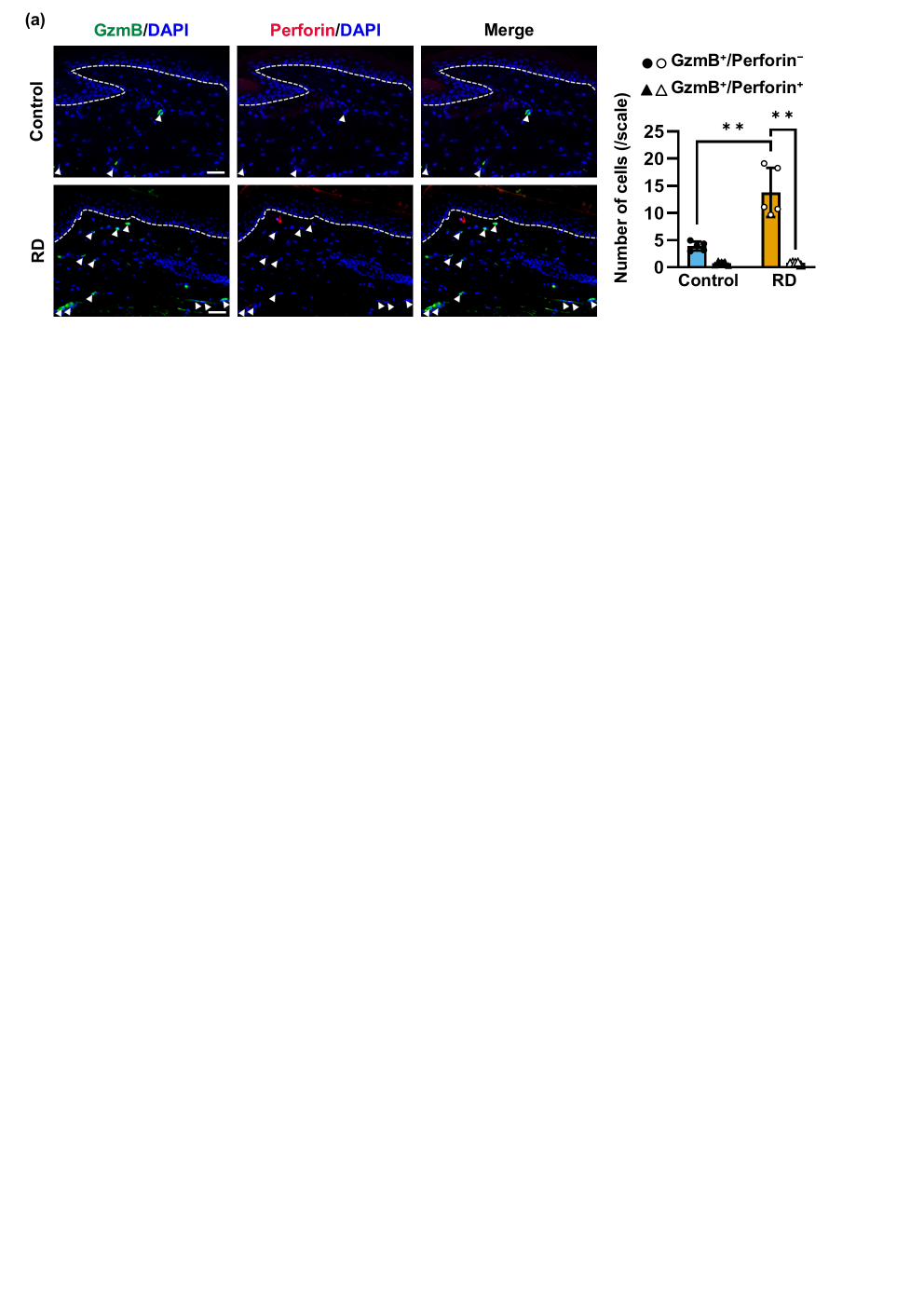
