## Supplementary figures and images for "Extracellular Granzyme B Promotes Melanocytorrhagy, Melanocyte Senescence and Aberrant Epidermal Differentiation in Vitiligo and Is Therapeutically Targetable"

### Fig S1a

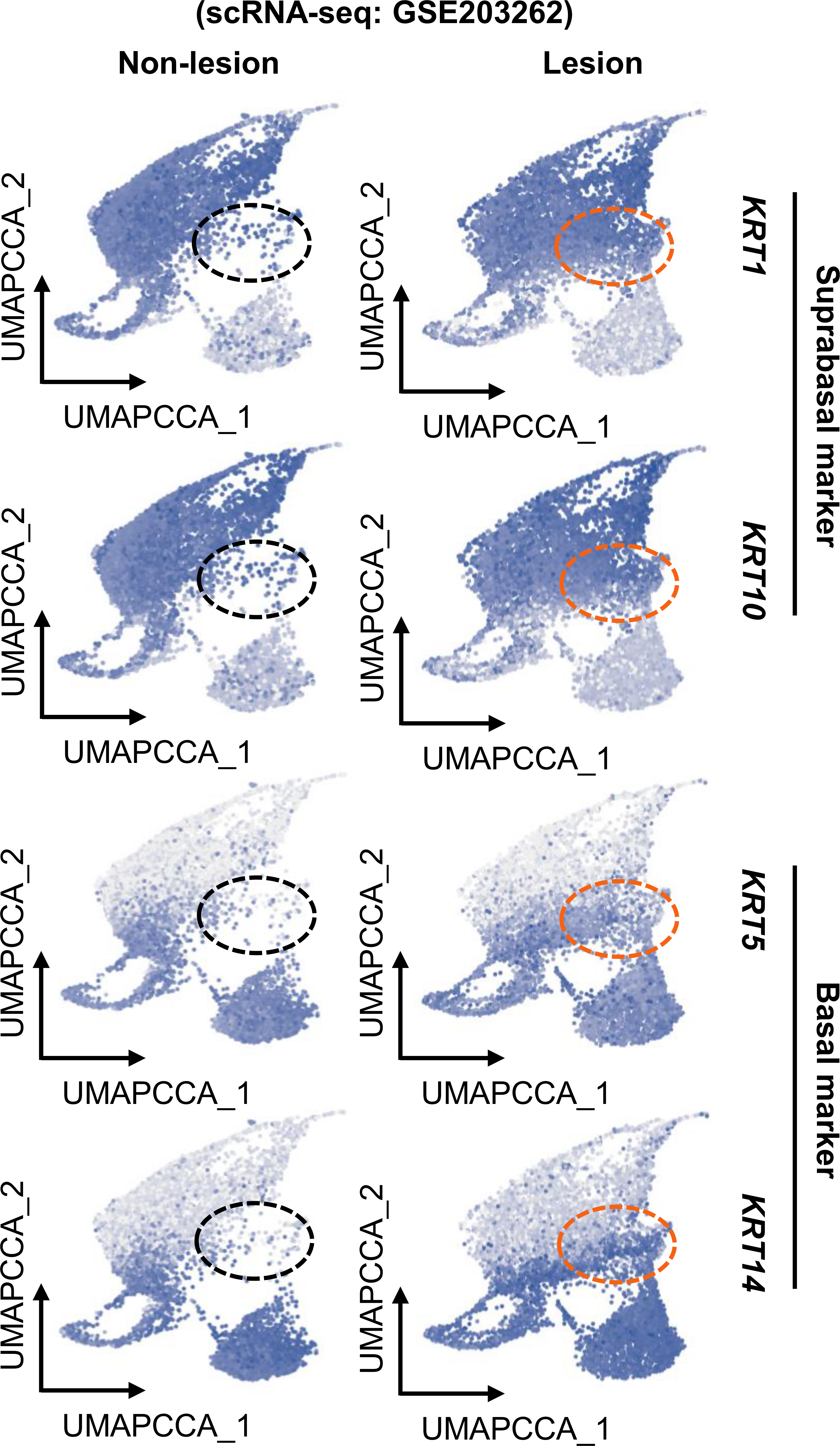

### Fig S1b

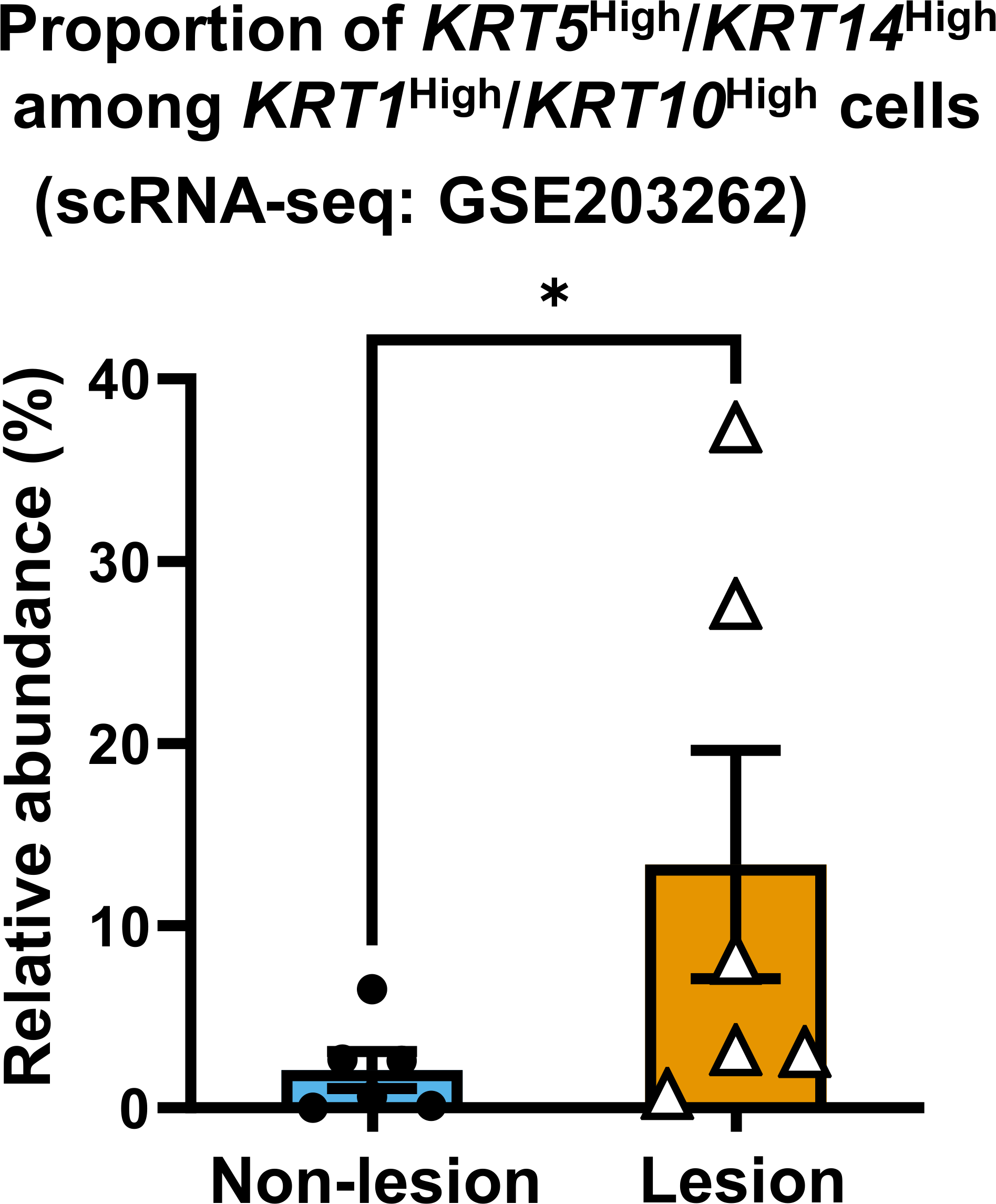

### Fig S2a

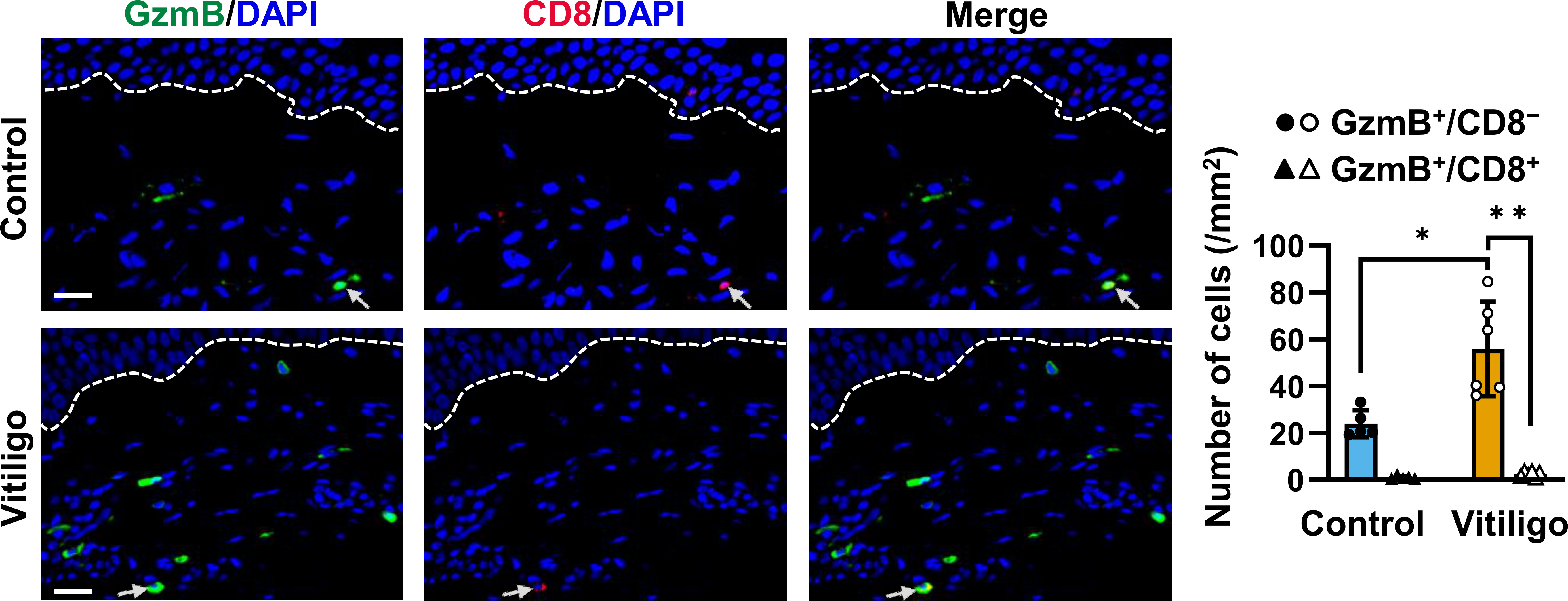

### Fig S2b

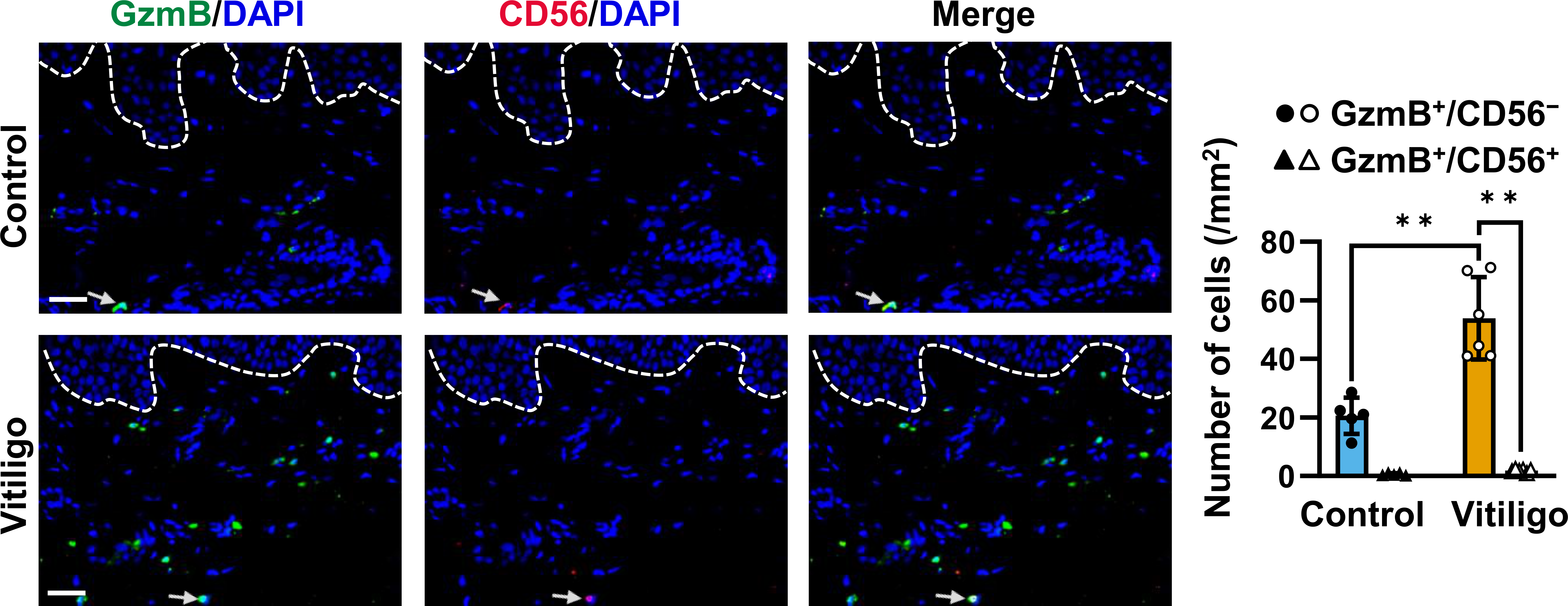

### Fig S2c

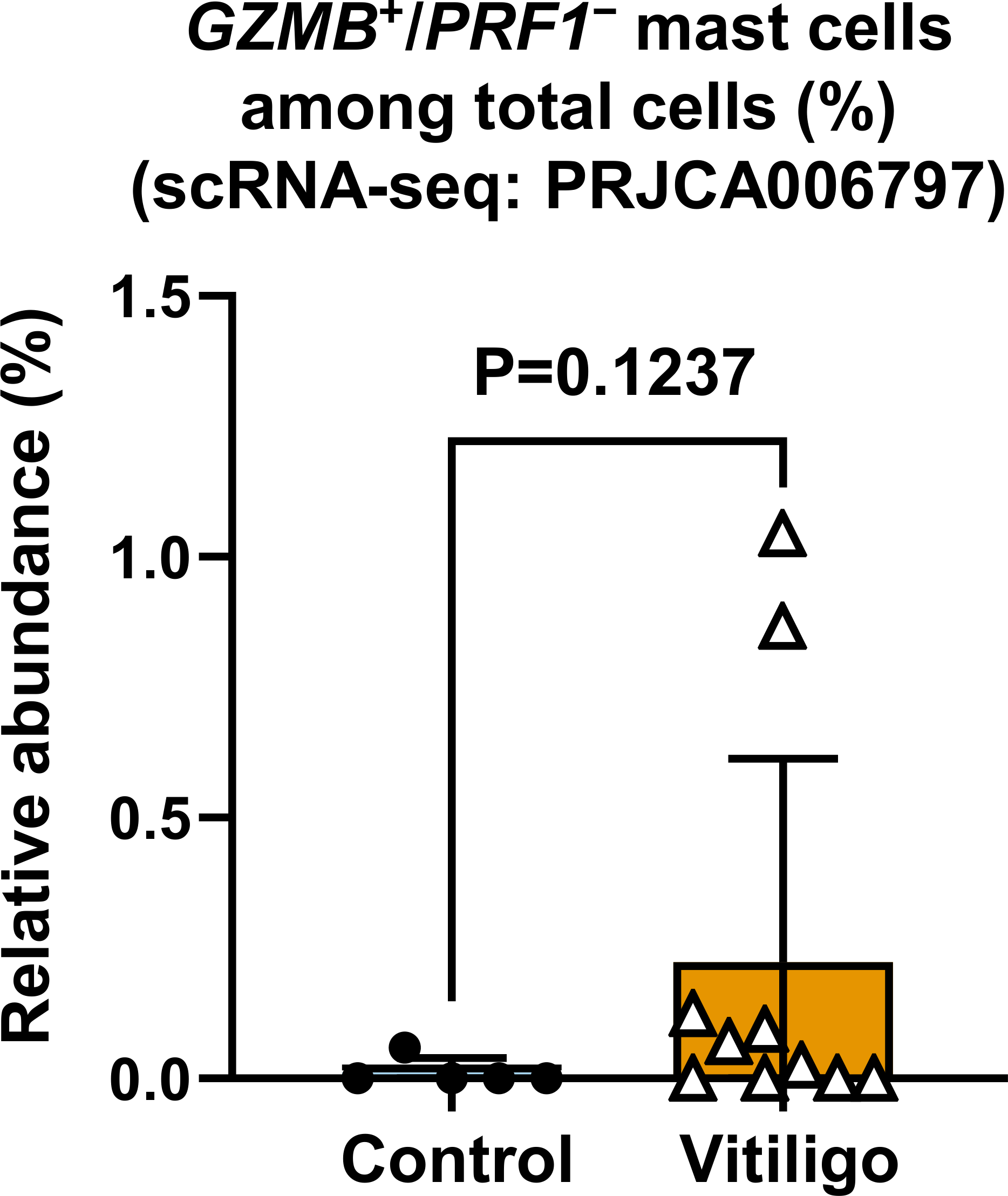

### Fig S3a

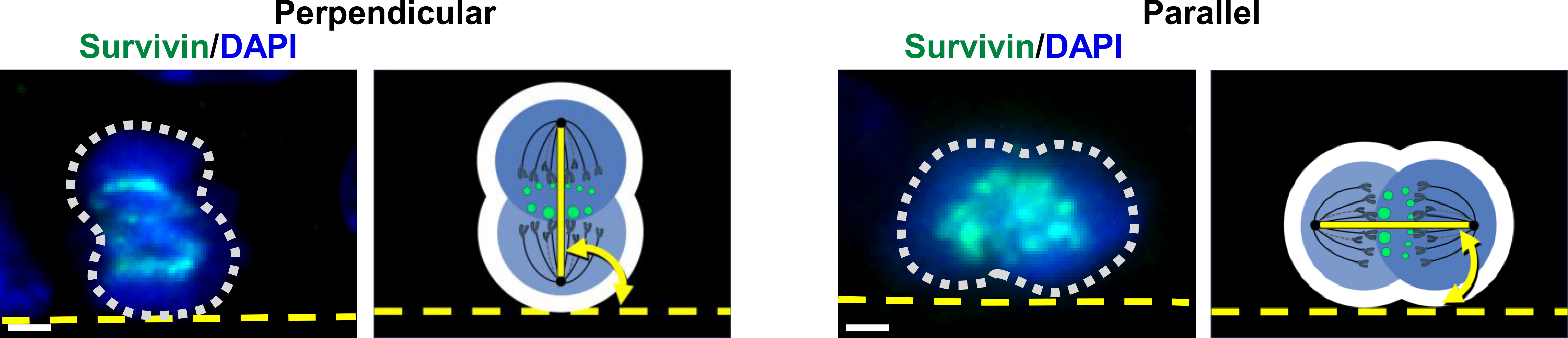

### Fig S3b

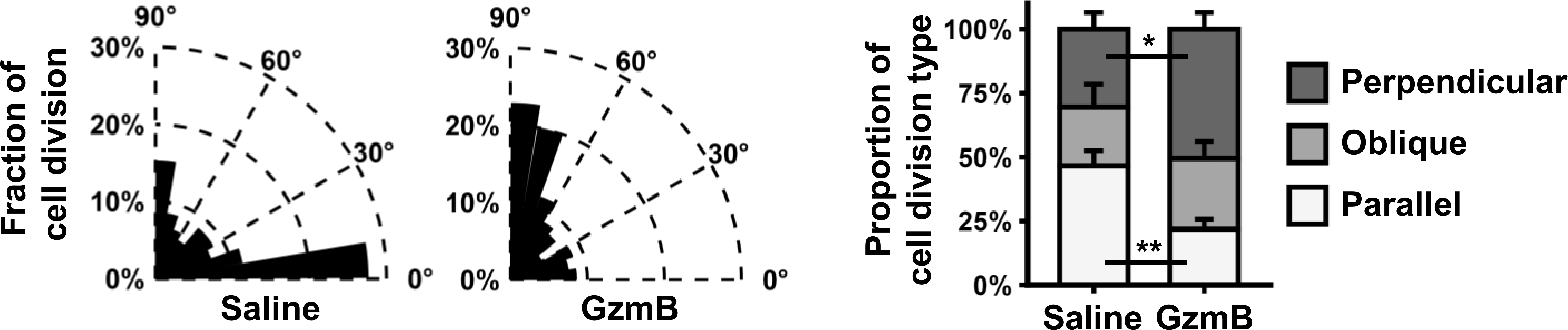

### Fig S3c

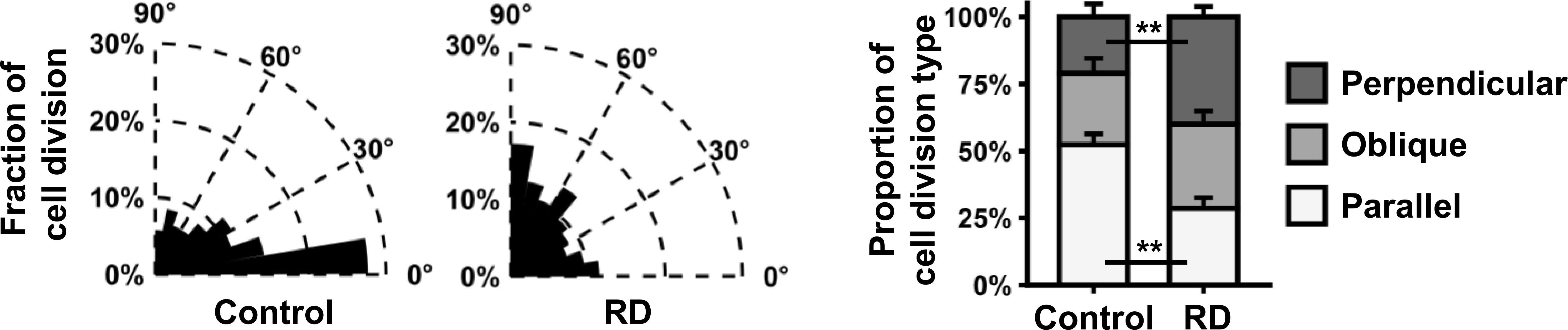

### Fig S3d

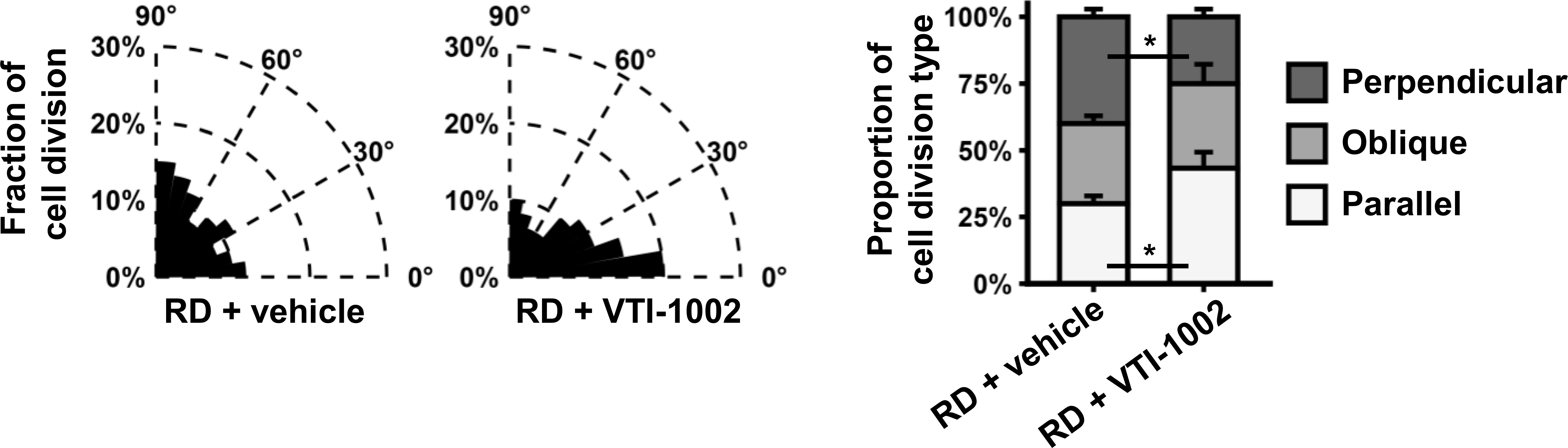

### Fig S4a

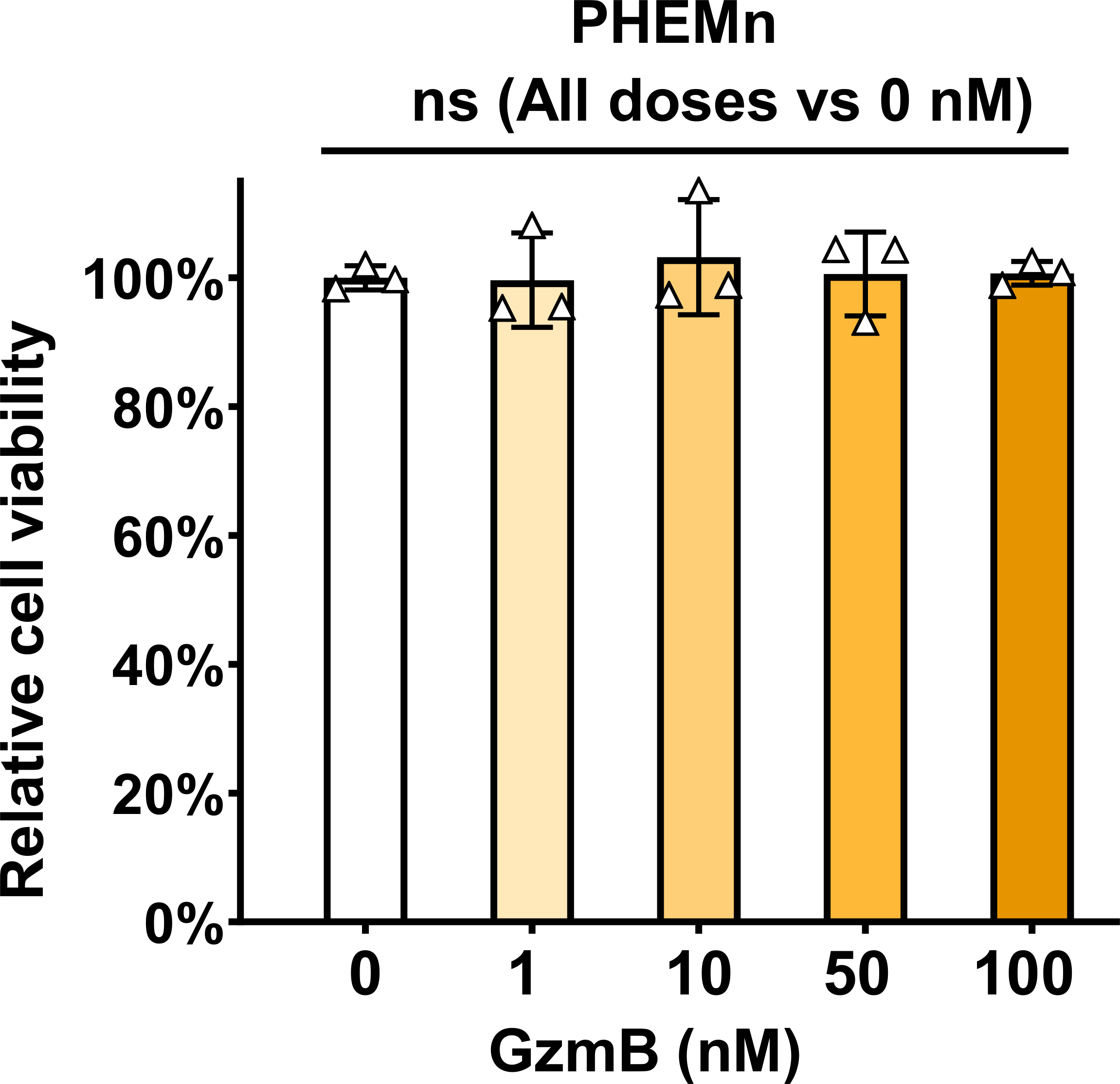

### Fig S4b

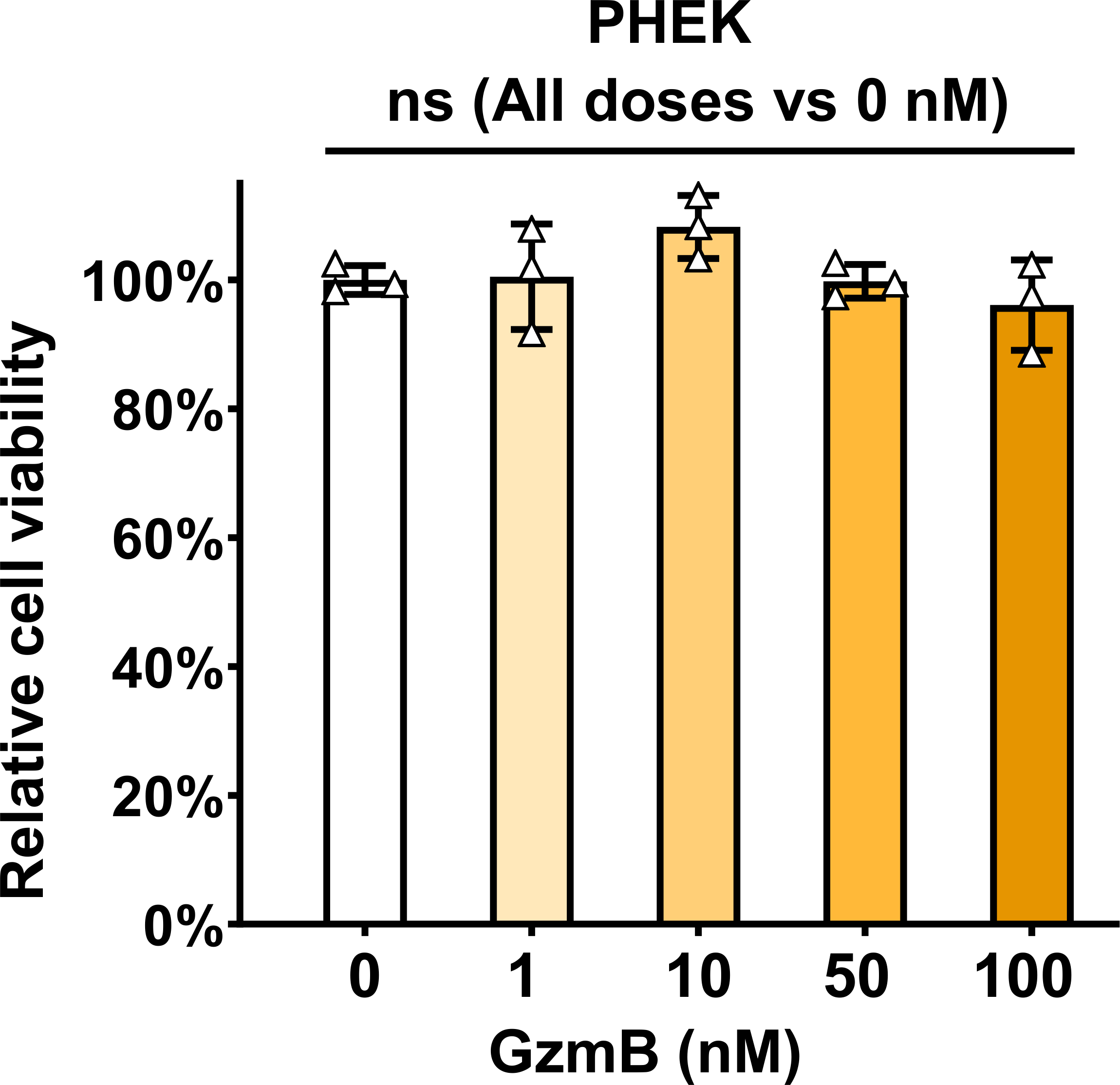

### Fig S5a

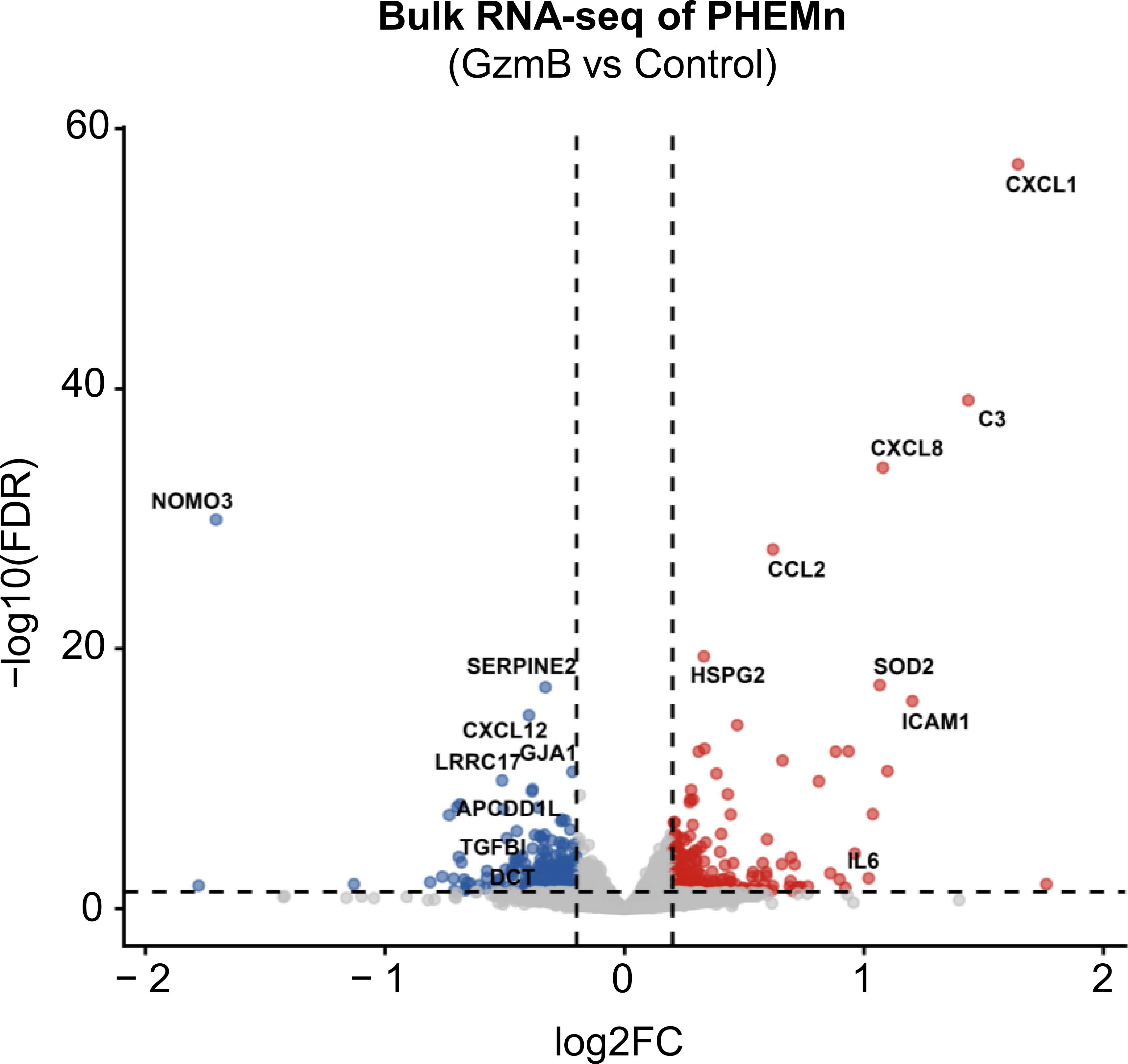

### Fig S5b

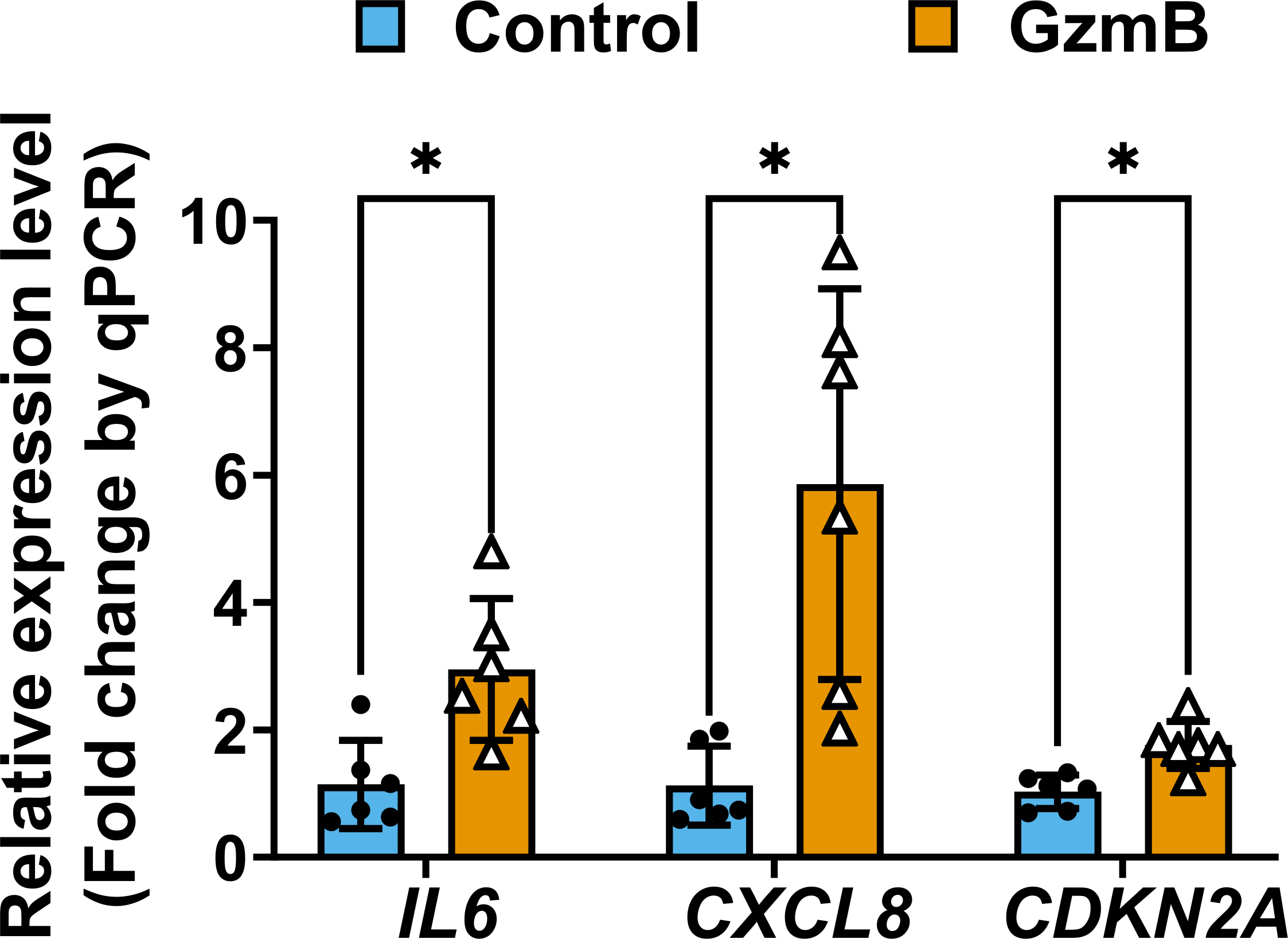

### Fig S5c

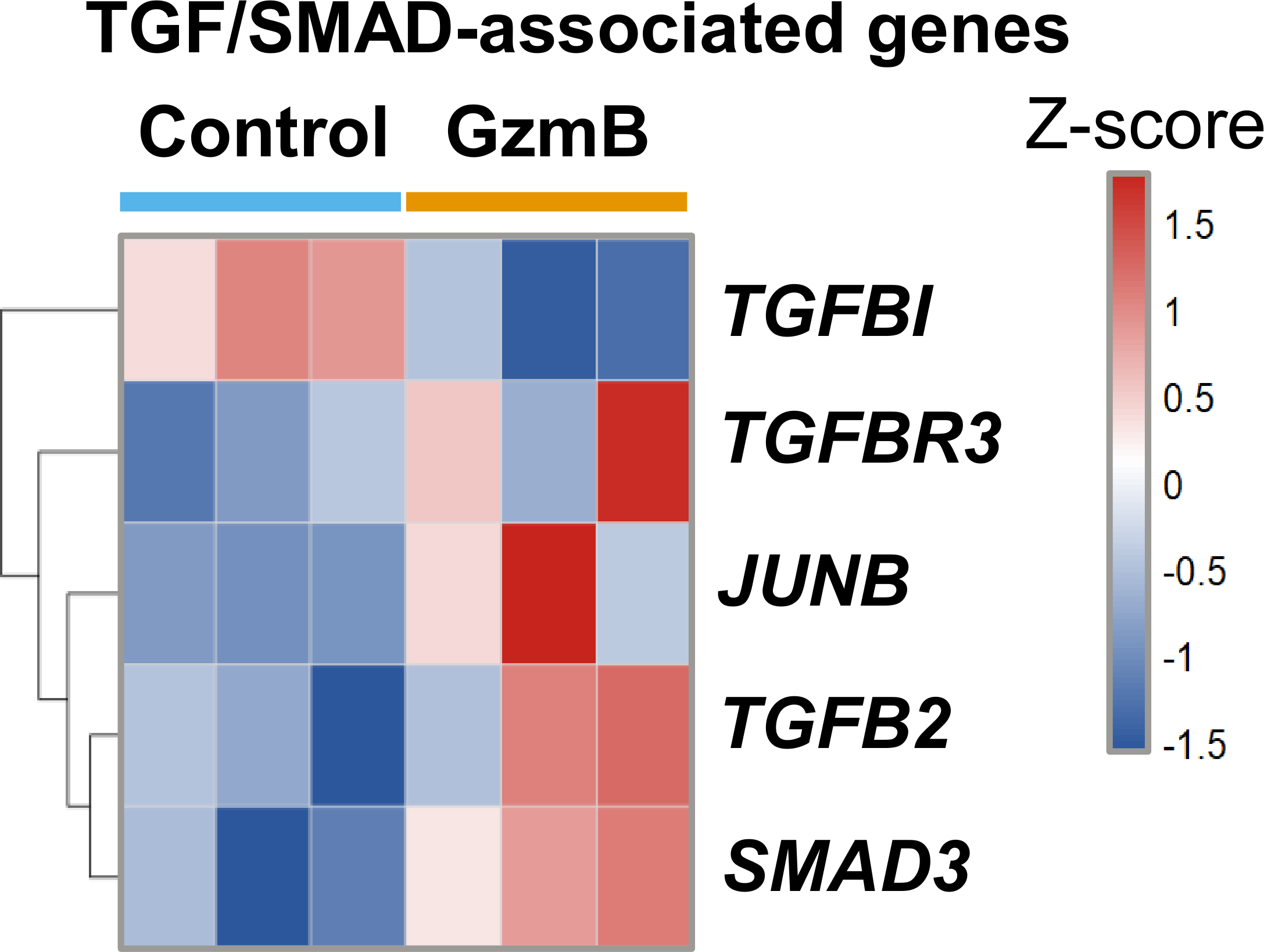

### Fig S5d

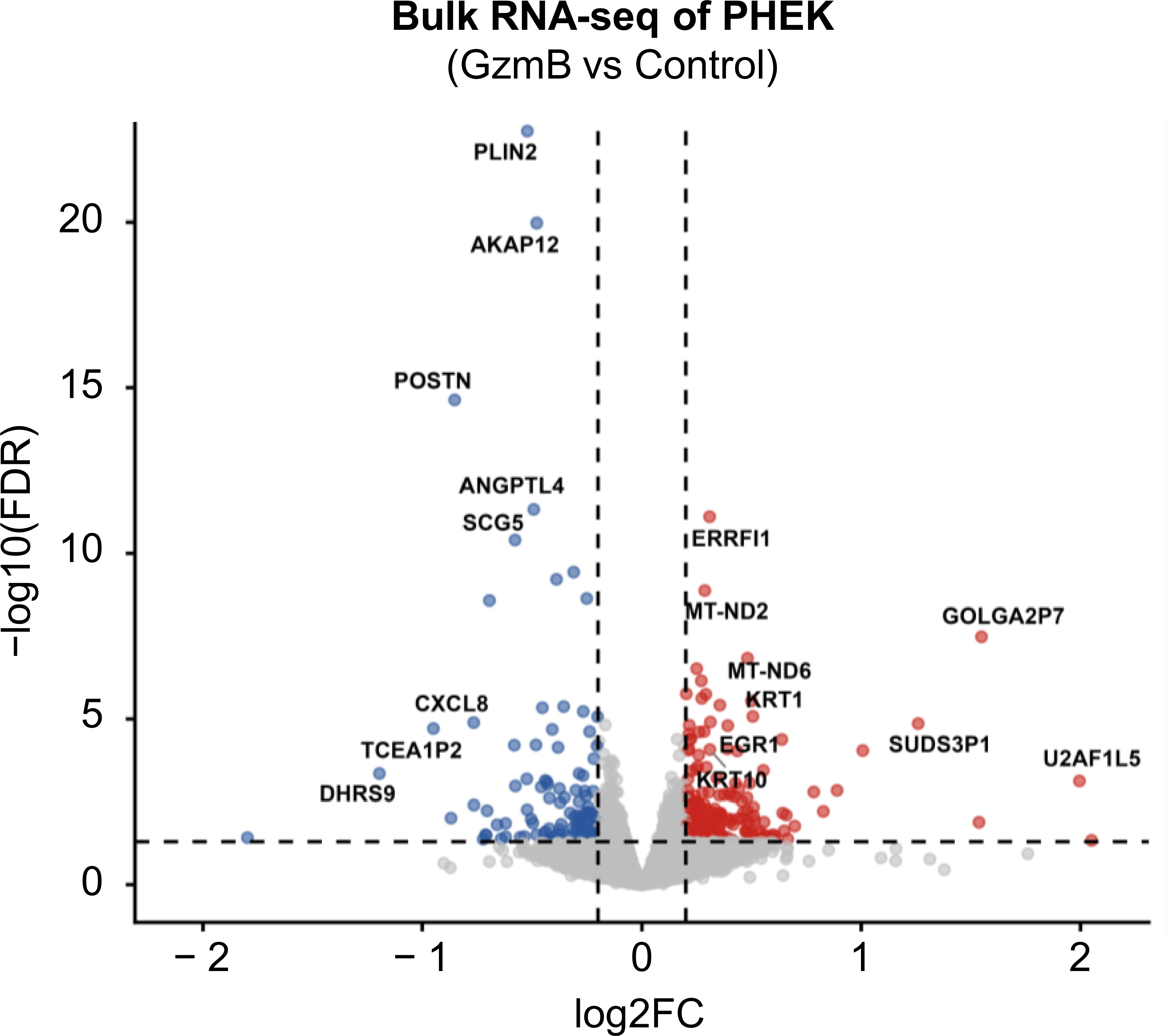

### Fig S5e

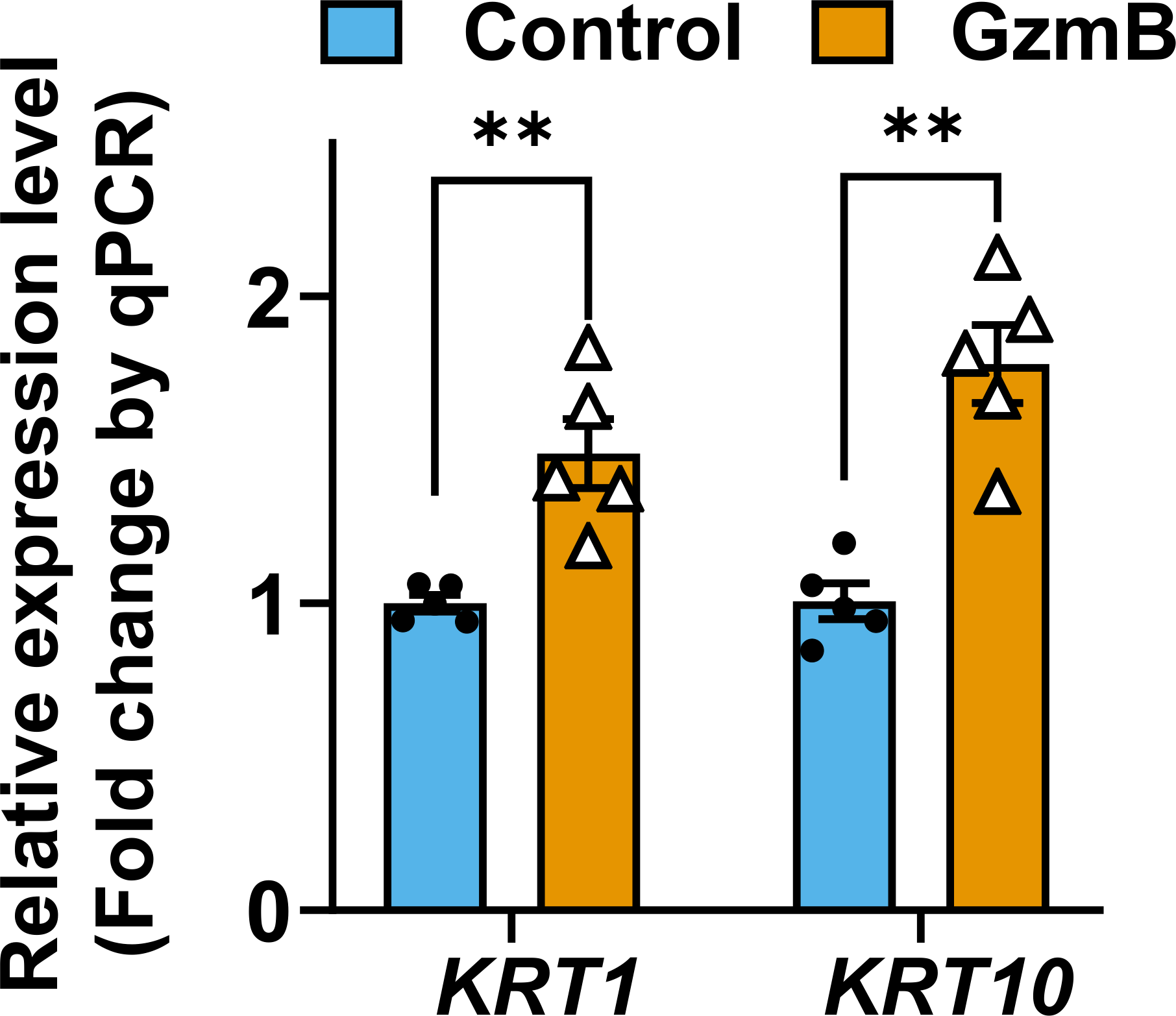

### Fig S5f

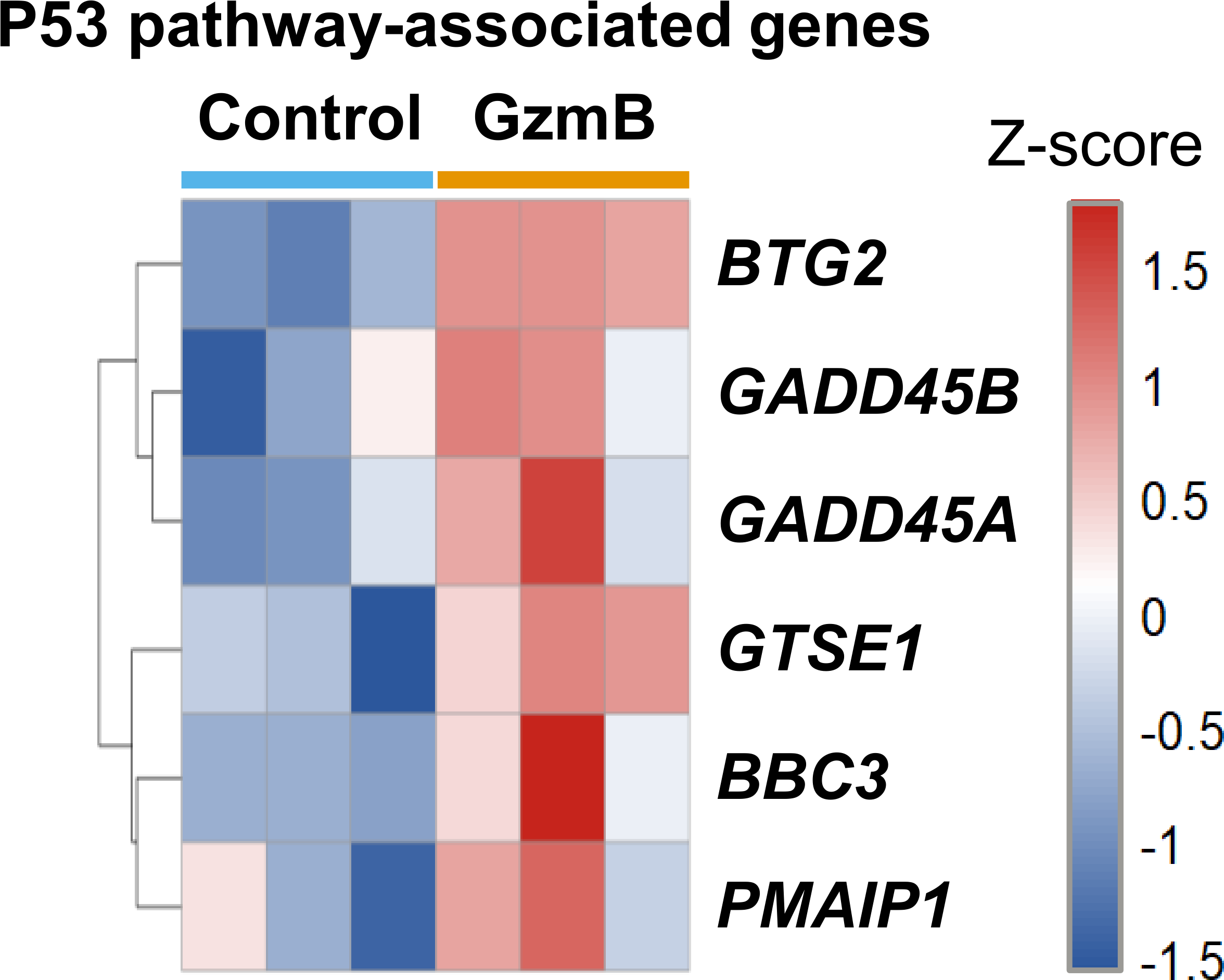

### Fig S6a

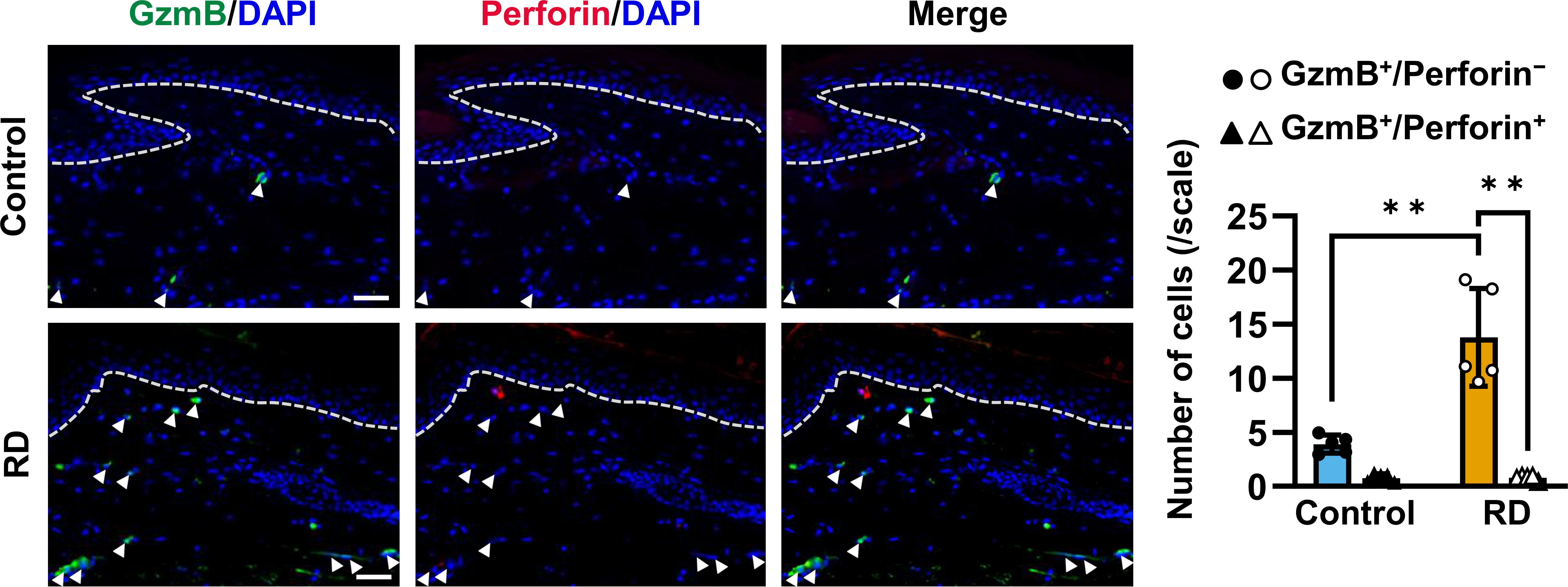
